# Variation in synonymous nucleotide composition among genomes of sarbecoviruses and consequences for the origin of COVID-19

**DOI:** 10.1101/2021.08.26.457807

**Authors:** Alexandre Hassanin

**Affiliations:** Institut de Systématique, Évolution, Biodiversité (ISYEB), Sorbonne Université, CNRS, EPHE, MNHN, UA, Paris

**Keywords:** synonymous mutations, coronavirus, *Manis javanica*, intermediate host, secondary host, reservoir host, hibernation

## Abstract

The subgenus *Sarbecovirus* includes two human viruses, SARS-CoV and SARS-CoV-2, respectively responsible for the SARS epidemic and COVID-19 pandemic, as well as many bat viruses and two pangolin viruses.

Here, the synonymous nucleotide composition (SNC) of *Sarbecovirus* genomes was analysed by examining third codon-positions, dinucleotides, and degenerate codons. The results show evidence for the eigth following groups: (i) SARS-CoV related coronaviruses (*SCoVrC* including many bat viruses from China), (ii) SARS-CoV-2 related coronaviruses (*SCoV2rC*; including five bat viruses from Cambodia, Thailand and Yunnan), (iii) pangolin viruses, (iv) three bat viruses showing evidence of recombination between *SCoVrC* and *SCoV2rC* genomes, (v) two highly divergent bat viruses from Yunnan, (vi) the bat virus from Japan, (vii) the bat virus from Bulgaria, and (viii) the bat virus from Kenya. All these groups can be diagnosed by specific nucleotide compositional features except the one concerned by recombination between *SCoVrC* and *SCoV2rC*. In particular, *SCoV2rC* genomes are characterised by the lowest percentages of cyosine and highest percentages of uracil at third codon-positions, whereas the genomes of pangolin viruses exhibit the highest percentages of adenine at third codon-positions. I suggest that latitudinal and taxonomic differences in the imbalanced nucleotide pools available in host cells during viral replication can explain the seven groups of SNC here detected among *Sarbecovirus* genomes. A related effect due to hibernating bats is also considered. I conclude that the two independent host switches from *Rhinolophus* bats to pangolins resulted in convergent mutational constraints and that SARS-CoV-2 emerged directly from a horseshoe bat virus.

## Introduction

The Severe Acute Respiratory Syndrome coronavirus 2 (SARS-CoV-2), which is the causative agent of the coronavirus disease 2019 (COVID-19), was first detected in December 2019 in Wuhan (China) (Wu et al. 2020). By the end of August 2021, the virus had spread to 220 countries and territories, causing more than 214 millions confirmed infections and 4.5 millions deaths (https://www.worldometers.info/coronavirus/). The SARS-CoV-2 is an enveloped virus containing a positive single-stranded RNA genome of 29.9 kb (Wu et al. 2020). After entry into the host cell, two large overlapping open reading frames (ORFs), ORF1a and ORF1b, are translated into polypeptides that are cleaved into 16 non-structural proteins involved in the viral replication and transcription complex. Then, the viral replication is initiated by the synthesis of full-length negative-sense genomic copies, which serve as templates for the synthesis of genomic and subgenomic RNAs (Finkel et al. 2021; V’kovski et al 2021). The different subgenomic RNAs encode the four coronavirus structural proteins - spike (S), envelope (E), membrane (M) and nucleocapsid (N) - and a number of accessory proteins (six were predicted in the reference SARS-CoV-2 genome NC_045512: ORF3a, ORF6, ORF7a, ORF7b, ORF8 and ORF10; Wu et al. 2020).

Phylogenetic analyses based on genomic sequences have shown that SARS-CoV-2 belongs to the subgenus *Sarbecovirus* of the family Coronaviridae, which also includes another human virus, SARS-CoV, involved in the SARS epidemic between 2002 and 2004, two pangolin viruses, and a large diversity of bat viruses (Boni et al. 2020; Lam et al. 2020; Zhou H. et al. 2020, 2021; Hul et al. 2021). Most of the bat sarbecoviruses were described from different species of the genus *Rhinolophus* (horseshoe bats) captured in caves of several provinces of China (Fan et al. 2019; Zhou H. et al. 2020, 2021). In addition, a few sarbecoviruses were detected in *Rhinolophus* species from Europe (Drexler et al. 2010), Africa (Tao and Tong 2019) and Southeast Asia (Hul et al. 2021; Wacharapluesadee et al. 2021), suggesting that horseshoe bats of the Old World constitute the reservoir host in which sarbecoviruses have been circulating and evolving for centuries.

Five SARS-CoV-2 related coronaviruses (*SCoV2rC*), sharing between 92 and 96% of genomic identity with SARS-CoV-2, were recently sequenced from five horseshoe bat species sampled in Yunnan (*Rhinolophus affinis*, *Rhinolophus malayanus*, and *Rhinolophus pusillus*), Cambodia (*Rhinolophus shameli*) and Thailand (*Rhinolophus acuminatus*) (Zhou H. et al. 2020; Zhou P. et al. 2020; Hul et al. 2021; Wacharapluesadee et al. 2021; Zhou et al. 2021). Based on these discoveries, the ecological niche of bat *SCoV2rC* was inferred using phylogeographic analyses of *Rhinolophus* species and it was found to include the four following geographic areas (Hassanin et al. 2021b): (i) southern Yunnan, northern Laos and bordering regions in northern Thailand and northwestern Vietnam; (ii) southern Laos, southwestern Vietnam, and northeastern Cambodia; (iii) the Cardamom Mountains in southwestern Cambodia and the East region of Thailand; and (iv) the Dawna Range in central Thailand and southeastern Myanmar. Importantly, the distribution of Sunda pangolin (*Manis javanica*) covers all these four geographic areas, as well as most other regions of mainland Southeast Asia, Borneo, Sumatra and Java (IUCN 2021). Since two sarbecoviruses related to *SCoV2rC* were sequenced from several Sunda pangolins seized in China between 2017 and 2019 (Lam et al. 2020; Xiao et al. 2020), it has been suggested that the species *Manis javanica* may have served as intermediate host between bat reservoirs and humans to import the ancestor of SARS-CoV-2 from Yunnan or Southeast Asia to the Chinese province of Hubei through wildlife trafficking (Lam et al. 2020; Hassanin et al. 2021a). In accordance with the hypothesis, pangolins are known to be highly permissive to contamination by sarbecoviruses (Xiao et al. 2020), as are also several small carnivores raised for fur or meat in China, such as the masked palm civet (*Paguma larvata*), raccoon dog (*Nyctereutes procyonoides*) (Guan et al. 2003) and American mink (*Neovison vison*) (Oude Munnink et al. 2021). In addition, it has been shown that pangolins collected in different geographic localities of Southeast Asia have been contaminated during their captivity in China (Hassanin et al. 2021a), reinforcing their possible role as intermediate host. However, all the closest relatives of SARS-CoV-2 currently known in wild animals were detected in horseshoe bats, not in pangolins or small carnivores, suggesting that the human index case was directly contaminated by a bat virus.

The sarbecoviruses are obligate intracellular pathogens that cannot replicate without the machinery of a host cell. After entrance into the host cell, the replication of their positive-strand RNA genome is initiated by the synthesis of full-length negative-sense genomic copies, which function as templates for the generation of new RNA genomes (V’kovski et al. 2021). Since the replication process is dependent of the host cell, a viral host-shift to a new mammalian species (i.e., from the reservoir to a secondary host or from an intermediate host to a terminal host) may result in important changes in the mutational patterns driving the evolution of viral genomes. For instance, it has been shown that most SARS-CoV-2 mutations in human populations are represented by C=>U transitions (Rice et al. 2020; Matyášek et al. 2021). With time, such a mutational bias can affect the nucleotide content of viral genomes. Variation in nucleotide composition is generally studied at synonymous sites – where all types of mutations (at four-fold degenerate sites) or some of them (only transitions at two-fold degenerate sites) do not alter the sequences of amino acids encoded by the genes – because their evolution is assumed to be neutral or nearly so, i.e., weakly affected by natural selection (Kimura 1968; Ohta 1992). In the present study, the synonymous nucleotide composition (SNC) was therefore analysed in a selection of 54 *Sarbecovirus* genomes to provide new insight on the issue of the intermediate host. The three main objectives were (i) to evidence potential differences among *Sarbecovirus* genomes, (ii) to test wether the genomes of the two divergent pangolin viruses have similar SNCs or not, and (iii) to determine if the genomic SNC of SARS-CoV-2 exhibits some unusual features or if it is close to that of related sarbecoviruses found in horseshoe bats and Sunda pangolins.

## Materials and Methods

### Genomic alignment of *Sarbecovirus* sequences

Full genomes of *Sarbecovirus* available in May 2021 in GenBank (https://www.ncbi.nlm.nih.gov/) and GISAID (https://www.epicov.org/) databases were donwloaded in Fasta format. Sequences with large stretch of missing data were removed. Only a single sequence was retained for similar genomes showing less than 0.1% of nucleotide divergence, such as those available for human SARS-CoV-2 (millions of sequences), pangolin viruses from Guangxi (5 sequences), bat viruses from Thailand (5 sequences), etc. All viral lineages previously described within the subgenus *Sarbecovirus* are included in this study (Drexler et al. 2010; Tao and Tong 2019; Murakami et al. 2020; Xiao et al. 2020; Zhou H. et al. 2020; Hul et al. 2021; Wacharapluesadee et al. 2021; Zhou et al. 2021). The details on the 54 selected genomes are provided in Table 1. The protein coding sequences (cds) of the 54 genomes were aligned in Geneious Prime® 2020.0.3 with MAFFT version 7.450 (Katoh and Standley 2013) using default parameters. Then, the alignment was corrected manually on AliView 1.26 (Larsson 2014) based on translated and untranslated nucleotide sequences using the three following criteria: (i) the number of indels was minimized because they are rarer events than amino-acid or nucleotide substitutions; (ii) changes between similar amino-acids were preferred; and (iii) transitions were privileged over transversions because they are more frequent. The final alignement contains 29,550 nucleotides (nt), representing 9,850 codons. It is available in the Open Science Framework platform at https://osf.io/XXXX/.

**Table 1.**
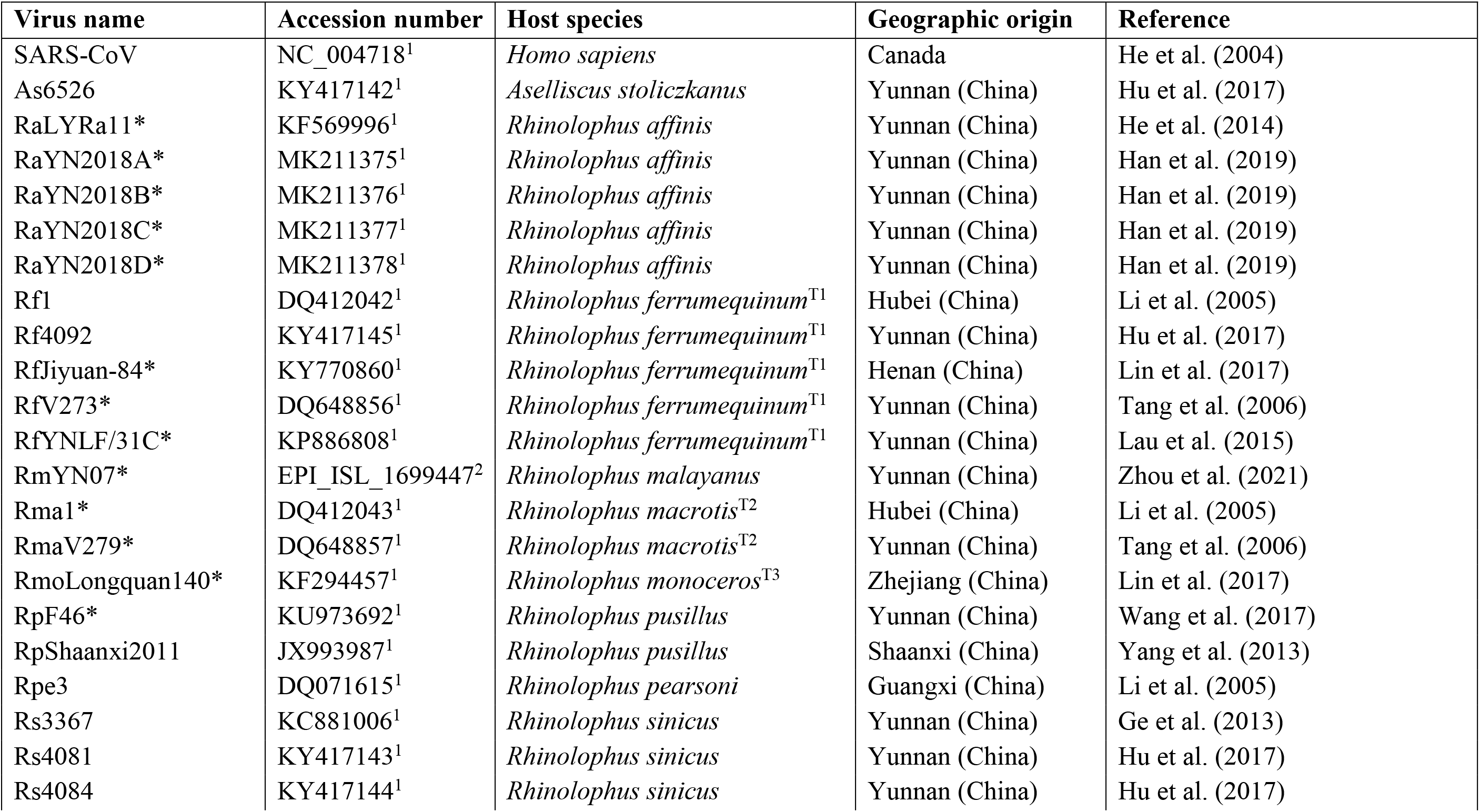
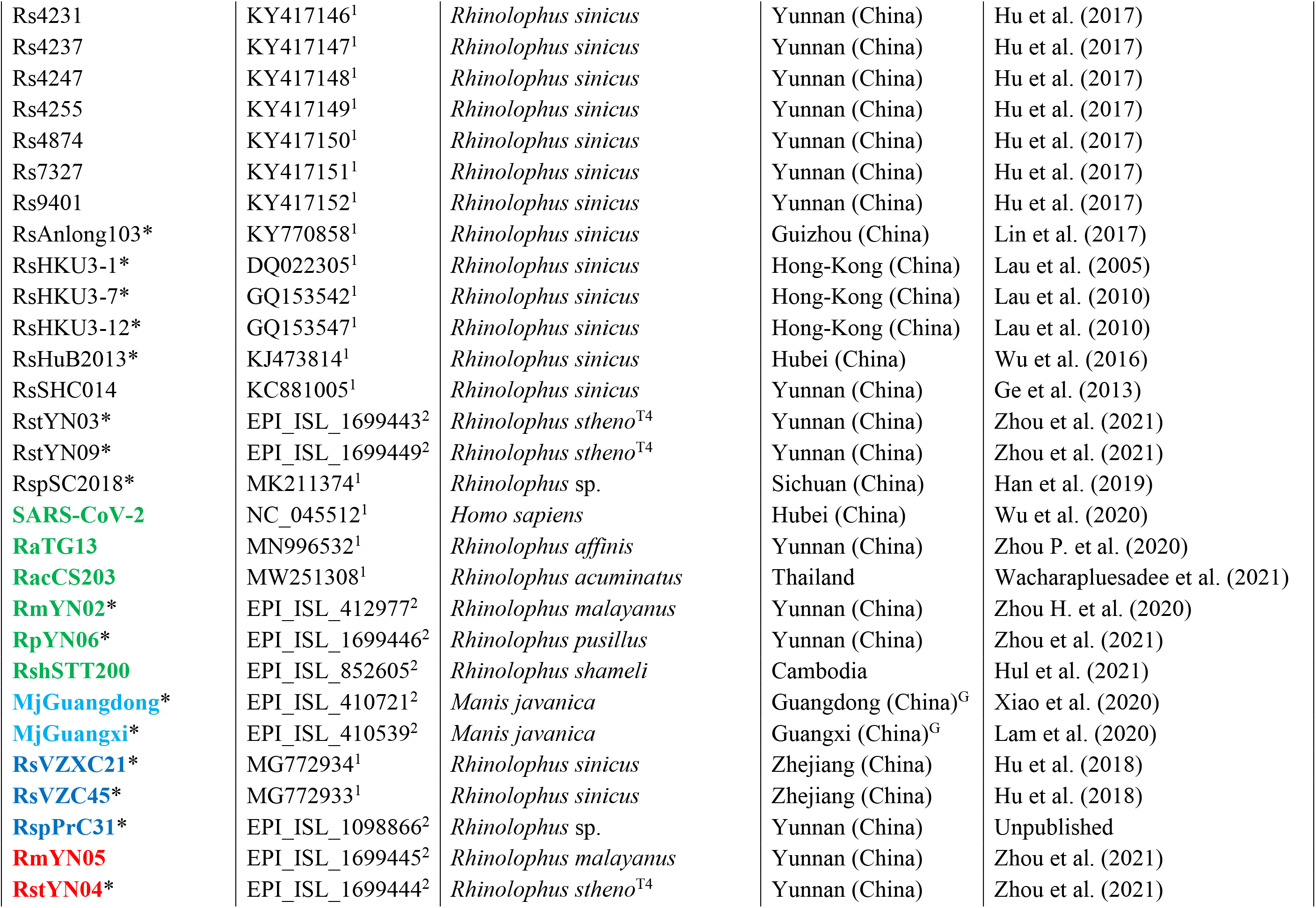
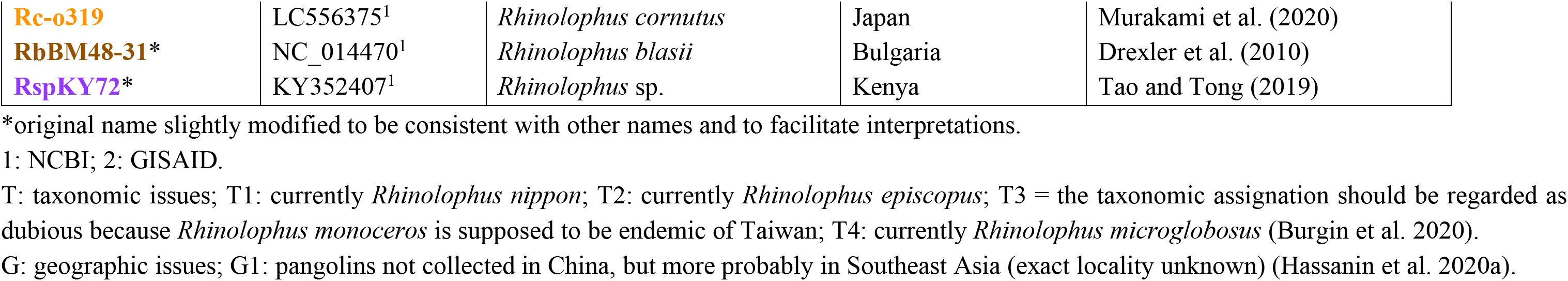
Origin of the *Sarbecovirus* genomes used in this study.

| Virus name | Accession number | Host species | Geographic origin | Reference |
| --- | --- | --- | --- | --- |
| SARS-CoV | NC_004718<sup>1</sup> | *Homo sapiens* | Canada | He et al. (2004) |
| As6526 | KY417142<sup>1</sup> | *Aselliscus stoliczkanus* | Yunnan (China) | Hu et al. (2017) |
| RaLYRa11* | KF569996<sup>1</sup> | *Rhinolophus affinis* | Yunnan (China) | He et al. (2014) |
| RaYN2018A* | MK211375<sup>1</sup> | *Rhinolophus affinis* | Yunnan (China) | Han et al. (2019) |
| RaYN2018B* | MK211376<sup>1</sup> | *Rhinolophus affinis* | Yunnan (China) | Han et al. (2019) |
| RaYN2018C* | MK211377<sup>1</sup> | *Rhinolophus affinis* | Yunnan (China) | Han et al. (2019) |
| RaYN2018D* | MK211378<sup>1</sup> | *Rhinolophus affinis* | Yunnan (China) | Han et al. (2019) |
| Rf1 | DQ412042<sup>1</sup> | *Rhinolophus ferrumequinum*<sup>T1</sup> | Hubei (China) | Li et al. (2005) |
| Rf4092 | KY417145<sup>1</sup> | *Rhinolophus ferrumequinum*<sup>T1</sup> | Yunnan (China) | Hu et al. (2017) |
| RfJiyuan-84* | KY770860<sup>1</sup> | *Rhinolophus ferrumequinum*<sup>T1</sup> | Henan (China) | Lin et al. (2017) |
| RfV273* | DQ648856<sup>1</sup> | *Rhinolophus ferrumequinum*<sup>T1</sup> | Yunnan (China) | Tang et al. (2006) |
| RfYNLF/31C* | KP886808<sup>1</sup> | *Rhinolophus ferrumequinum*<sup>T1</sup> | Yunnan (China) | Lau et al. (2015) |
| RmYN07* | EPI_ISL_1699447<sup>2</sup> | *Rhinolophus malayanus* | Yunnan (China) | Zhou et al. (2021) |
| Rma1* | DQ412043<sup>1</sup> | *Rhinolophus macrotis*<sup>T2</sup> | Hubei (China) | Li et al. (2005) |
| RmaV279* | DQ648857<sup>1</sup> | *Rhinolophus macrotis*<sup>T2</sup> | Yunnan (China) | Tang et al. (2006) |
| RmoLongquan140* | KF294457<sup>1</sup> | *Rhinolophus monoceros*<sup>T3</sup> | Zhejiang (China) | Lin et al. (2017) |
| RpF46* | KU973692<sup>1</sup> | *Rhinolophus pusillus* | Yunnan (China) | Wang et al. (2017) |
| RpShaanxi2011 | JX993987<sup>1</sup> | *Rhinolophus pusillus* | Shaanxi (China) | Yang et al. (2013) |
| Rpe3 | DQ071615<sup>1</sup> | *Rhinolophus pearsoni* | Guangxi (China) | Li et al. (2005) |
| Rs3367 | KC881006<sup>1</sup> | *Rhinolophus sinicus* | Yunnan (China) | Ge et al. (2013) |
| Rs4081 | KY417143<sup>1</sup> | *Rhinolophus sinicus* | Yunnan (China) | Hu et al. (2017) |
| Rs4084 | KY417144<sup>1</sup> | *Rhinolophus sinicus* | Yunnan (China) | Hu et al. (2017) |
| Rs4231 | KY417146<sup>1</sup> | *Rhinolophus sinicus* | Yunnan (China) | Hu et al. (2017) |
| Rs4237 | KY417147<sup>1</sup> | *Rhinolophus sinicus* | Yunnan (China) | Hu et al. (2017) |
| Rs4247 | KY417148<sup>1</sup> | *Rhinolophus sinicus* | Yunnan (China) | Hu et al. (2017) |
| Rs4255 | KY417149<sup>1</sup> | *Rhinolophus sinicus* | Yunnan (China) | Hu et al. (2017) |
| Rs4874 | KY417150<sup>1</sup> | *Rhinolophus sinicus* | Yunnan (China) | Hu et al. (2017) |
| Rs7327 | KY417151<sup>1</sup> | *Rhinolophus sinicus* | Yunnan (China) | Hu et al. (2017) |
| Rs9401 | KY417152<sup>1</sup> | *Rhinolophus sinicus* | Yunnan (China) | Hu et al. (2017) |
| RsAnlong103* | KY770858<sup>1</sup> | *Rhinolophus sinicus* | Guizhou (China) | Lin et al. (2017) |
| RsHKU3-1* | DQ022305<sup>1</sup> | *Rhinolophus sinicus* | Hong-Kong (China) | Lau et al. (2005) |
| RsHKU3-7* | GQ153542<sup>1</sup> | *Rhinolophus sinicus* | Hong-Kong (China) | Lau et al. (2010) |
| RsHKU3-12* | GQ153547<sup>1</sup> | *Rhinolophus sinicus* | Hong-Kong (China) | Lau et al. (2010) |
| RsHuB2013* | KJ473814<sup>1</sup> | *Rhinolophus sinicus* | Hubei (China) | Wu et al. (2016) |
| RsSHC014 | KC881005<sup>1</sup> | *Rhinolophus sinicus* | Yunnan (China) | Ge et al. (2013) |
| RstYN03* | EPI_ISL_1699443<sup>2</sup> | *Rhinolophus stheno*<sup>T4</sup> | Yunnan (China) | Zhou et al. (2021) |
| RstYN09* | EPI_ISL_1699449<sup>2</sup> | *Rhinolophus stheno*<sup>T4</sup> | Yunnan (China) | Zhou et al. (2021) |
| RspSC2018* | MK211374<sup>1</sup> | *Rhinolophus* sp. | Sichuan (China) | Han et al. (2019) |
| **SARS-CoV-2** | NC_045512<sup>1</sup> | *Homo sapiens* | Hubei (China) | Wu et al. (2020) |
| **RaTG13** | MN996532<sup>1</sup> | *Rhinolophus affinis* | Yunnan (China) | Zhou P. et al. (2020) |
| **RacCS203** | MW251308<sup>1</sup> | *Rhinolophus acuminatus* | Thailand | Wacharapluesadee et al. (2021) |
| **RmYN02\*** | EPI_ISL_412977<sup>2</sup> | *Rhinolophus malayanus* | Yunnan (China) | Zhou H. et al. (2020) |
| **RpYN06\*** | EPI_ISL_1699446<sup>2</sup> | *Rhinolophus pusillus* | Yunnan (China) | Zhou et al. (2021) |
| **RshSTT200** | EPI_ISL_852605<sup>2</sup> | *Rhinolophus shameli* | Cambodia | Hul et al. (2021) |
| **MjGuangdong\*** | EPI_ISL_410721<sup>2</sup> | *Manis javanica* | Guangdong (China)<sup>G</sup> | Xiao et al. (2020) |
| **MjGuangxi\*** | EPI_ISL_410539<sup>2</sup> | *Manis javanica* | Guangxi (China)<sup>G</sup> | Lam et al. (2020) |
| **RsVZXC21\*** | MG772934<sup>1</sup> | *Rhinolophus sinicus* | Zhejiang (China) | Hu et al. (2018) |
| **RsVZC45\*** | MG772933<sup>1</sup> | *Rhinolophus sinicus* | Zhejiang (China) | Hu et al. (2018) |
| **RspPrC31\*** | EPI_ISL_1098866<sup>2</sup> | *Rhinolophus* sp. | Yunnan (China) | Unpublished |
| **RmYN05** | EPI_ISL_1699445<sup>2</sup> | *Rhinolophus malayanus* | Yunnan (China) | Zhou et al. (2021) |
| **RstYN04\*** | EPI_ISL_1699444<sup>2</sup> | *Rhinolophus stheno*<sup>T4</sup> | Yunnan (China) | Zhou et al. (2021) |
| **Rc-o319** | LC556375<sup>1</sup> | *Rhinolophus cornutus* | Japan | Murakami et al. (2020) |
| **RbBM48-31\*** | NC_014470<sup>1</sup> | *Rhinolophus blasii* | Bulgaria | Drexler et al. (2010) |
| **RspKY72\*** | KY352407<sup>1</sup> | *Rhinolophus* sp. | Kenya | Tao and Tong (2019) |

\*original name slightly modified to be consistent with other names and to facilitate interpretations.
1: NCBI; 2: GISAID.
T: taxonomic issues; T1: currently *Rhinolophus nippon*; T2: currently *Rhinolophus episcopus*; T3 = the taxonomic assignation should be regarded as dubious because *Rhinolophus monoceros* is supposed to be endemic of Taiwan; T4: currently *Rhinolophus microglobosus* (Burgin et al. 2020).
G: geographic issues; G1: pangolins not collected in China, but more probably in Southeast Asia (exact locality unknown) (Hassanin et al. 2020a).

### Phylogeny of sarbecoviruses

All phylogenetic analyses were carried out using the maximum likelihood (ML) method under RAxML version 8.2.10 (Stamatakis 2014) with 25 rate categories, different GTRCAT models for the three codon-positions, and 1000 bootstrap replicates. The 1000 bootstrap RAxML trees were then executed in PAUP 4* version 4b10 (Swofford 2003) to construct a 70% majority-rule consensus tree.

Phylogenetic relationships were first inferred using the whole genomic alignment of 29,550 nt. Since several studies have shown evidence for multiple events of genomic recombination during the evolutionary history of sarbecoviruses (Hon et al. 2008; Boni et al. 2020), phylogenetic analyses were also conducted using the three following cds regions of the alignment: the ORF1ab (positions 1-21429; 7143 codons), the RdRp gene (positions 13321-1613; 931 codons), and the Spike gene (positions 21433-25329; 1299 codons).

### Analyses of nucleotide composition, dinucleotide composition and codon usage

The alignment of 54 *Sarbecovirus* genomes was used to calculate the frequency of the four nucleotides (A, C, G and U) at all third codon-positions. Nucleotide frequencies were calculated in PAUP (Swofford 2003) after exclusion of first and second codon-positions. The four variables measured were then summarised by a principal component analysis (PCA) using the FactoMineR package (Lê et al. 2008) in R version 3.6.1 (from http://www.R-project.org/). The nucleotide composition was also determined using three partitions of third codon-positions consisting in the four-fold degenerate sites (A, C, G and U percentages), purine two-fold degenerate sites (A *versus* G percentages), and pyrimidine two-fold degenerate sites (C *versus* U percentages). The eight variables were summarised by a PCA.

The genomic alignment was also used to calculate the frequency of the 16 possible dinucleotides (AA, AC, AG, AU, CA, CC, CG, CU, GA, GC, GG, GU, UA, UC, UG, and UU) at second and third codon-positions, and at third and first codon-positions. For each of the two analyses, the 16 variables were summarised by a PCA.

Finally, the frequencies of synonymous codons (codon usage) were calculated for all amino acids except M and W, which are encoded by a single codon each (AUG and UGG, respectively). The 59 variables were summarised by a PCA.

## Results

### Phylogenetic relationships within the subgenus *Sarbecovirus*

The bootstrap 70% majority-rule consensus tree derived from the RAxML analysis of the whole alignment of protein-coding sequences (29,550 nt) is shown in Figure 1A, with bootstrap proportions (BP) indicated at the nodes. Three groups of interest were found to be monophyletic: (i) *SCoVrC* (BP = 100); (ii) a group here referred to as *RecSar* (BP = 83), which includes three bat viruses showing evidence of genomic recombination between *SCoV2rC* and *SCoVrC* (RspPrC31, RsVZC45, and RsVZXC21; see below for more details); and (iii) a group here referred to as *YunSar* (BP = 100), which is composed of two divergent bat viruses from Yunnan, i.e. RmYN05 and RstYN04. By contrast, *SCoV2rC* was found to be paraphyletic due to the inclusive placement of *RecSar* (BP = 83). The two pangolin viruses did not group together: MjGuangdong was found to be closely related to *RecSar* + *SCoV2rC* (BP = 83), whereas MjGuangxi appeared more distantly related.

**Figure 1.**
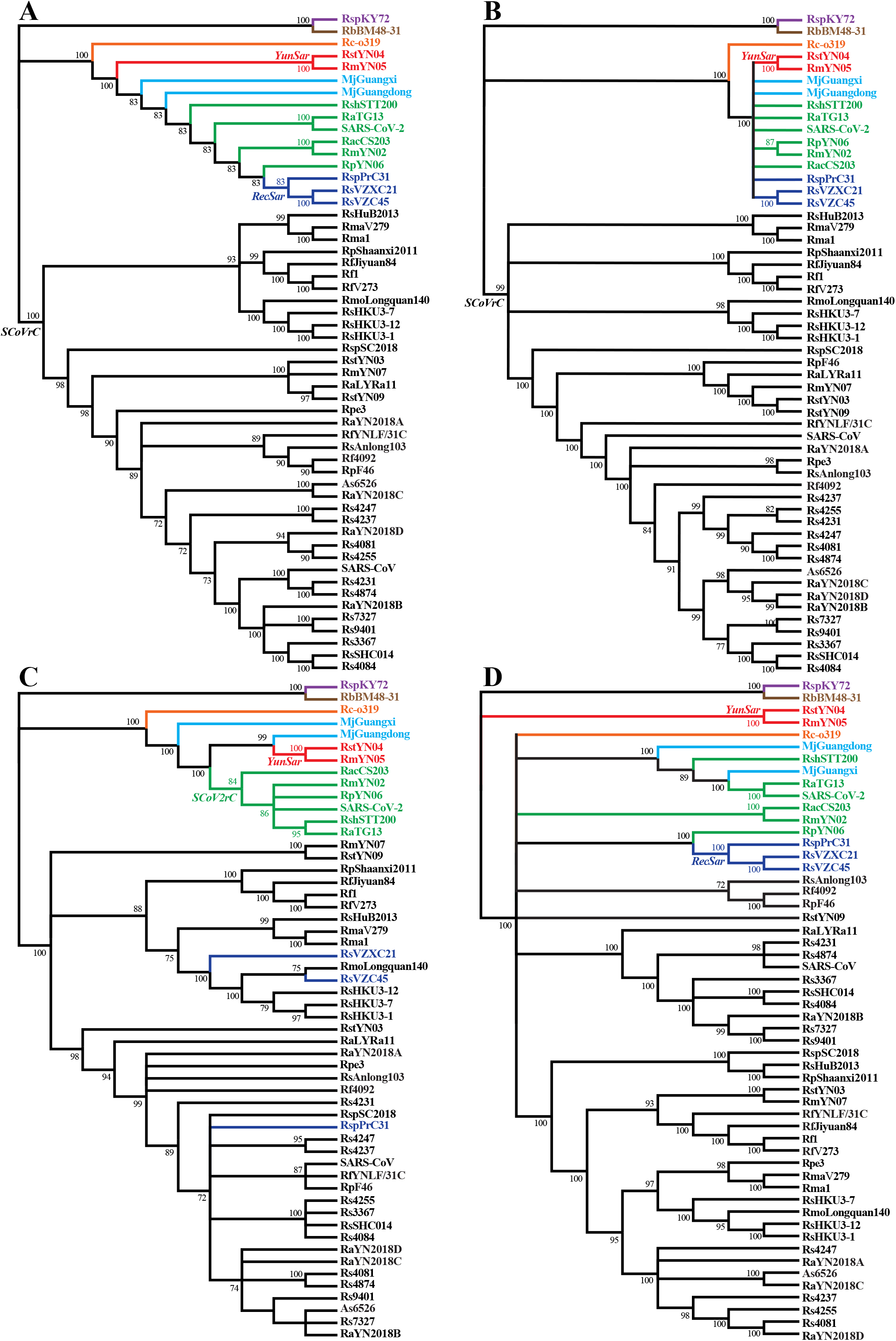
Phylogenetic relationships within the subgenus *Sarbecovirus* based on (A) the whole genomic alignment, (B) the ORF1ab, (C) the RdRp gene, and (D) the Spike gene. Phylogenetic analyses were performed with RAxML version 8.2.10 (Stamatakis 2014) using different GTRCAT models for the three codon-positions, 25 rate categories, and 1000 bootstrap replicates. The bootstrap 70% majority-rule consensus trees were constructed in PAUP 4* version 4b10 (Swofford 2003). All trees were rooted using RspKY72 and RbBM48-31 as outgroup. The colours indicate to which group of synonymous nucleotide composition (SNC) the viruses belong: black for coronaviruses related to SARS-CoV (SCoVrC), green for coronaviruses related to SARS-CoV-2 (SCoV2rC), light blue for the two pangolin viruses (*PangSar*), dark blue for bat viruses showing evidence of genomic recombination between SCoVrC and SCoV2rC (*RecSar*), red for the two divergent bat viruses recently identified in the Yunnan province by Zhou et al. (2021) (*YunSar*), orange for the bat virus from Japan, brown for the bat virus from Bulgaria, and purple for the bat virus from Kenya.

The bootstrap 70% majority-rule consensus tree constructed from the RAxML analysis of the ORF1ab (21,429 nt) is shown in Figure 1B. The tree was less resolved than the tree derived from the whole genomic alignment but deep relationships were found to be congruent, such as the monophyly of *SCoVrC* (BP = 99), the clade uniting *SCoV2rC*, *RecSar*, *YunSar*, and the two pangolin viruses (BP = 100), and its sister-group relationship with Rc-o319 (BP = 100). However, several more recent relationships were discordant. For instance, RmYN02 was closely related to RpYN06 in the ORF1ab tree (BP = 87), whereas it was the sister-group of RacCS203 in the whole genome tree (BP = 100).

The bootstrap 70% majority-rule consensus tree built from the RAxML analysis of the RdRp gene (2,793 nt) is shown in Figure 1C. The topology was highly incongruent with the tree inferred from the whole genomic alignment because the three *RecSar* viruses appeared included into the *SCoVrC* group: RsVZC45 was closely related to RmLongquan140 and three bat viruses from Hong-Kong (RsHKU3-1, -7, and -12) (BP = 100); RsVZXC21 was the sister-group of this clade (BP = 100); whereas RspPrC31 was enclosed into a large group including SARS-CoV and several bat viruses (BP = 98). In the RdRp tree, *SCoV2rC* was found to be monophyletic (BP = 84) and *YunSar* was closely related to MjGuangdong (BP = 99).

The bootstrap 70% majority-rule consensus tree derived from the RAxML analysis of the Spike gene (3,897 nt) is shown in Figure 1D. The tree was poorly resolved for deep relationships, but the group uniting *SCoVrC*, *SCoV2rC*, *RecSar*, Rc-o319 and the two pangolin viruses was found to be robust (BP = 100) due to the divergent placement of *YunSar*. The *SCoV2rC* goup was found to be polyphyletic: RpYN06 appeared closely related to *RecSar* (BP = 100), whereas RshSTT200 was allied to SARS-CoV-2 + RaTG13 (BP = 100) and MjGuangxi (BP = 100).

### Nucleotide composition at third codon-positions

The four nucleotide frequencies at third codon-positions are provided in Table 2 for the main groups identified in this study (CSV file available at https://osf.io/XXXX/). The four variables were summarised by a PCA based on the first two principal components (PCs), which contribute 89.08% and 9.84% of the total variance, respectively (Figure 2A). The results allowed to distinguish the eight following groups: (i) *SCoVrC*; (ii) *SCoV2rC*; (iii) pangolin sarbecoviruses (here referred to as *PangSar*); (iv) *RecSar*, including the three bat viruses showing evidence of recombination between *SCoV2rC* and *SCoVrC* (RspPrC31, RsVZXC21, and RsVZC45; see discussion for more details); (v) *YunSar*, including two highly divergent bat viruses from Yunnan (RmYN05 and RstYN04); (vi) the bat virus from Japan (Rc-o319); (vii) the bat virus from Bulgaria (RbBM48-31); and (viii) the bat virus from Kenya (RspKY72). As indicated in Table 2, *SCoV2rC* genomes are characterised by more U nucleotides and less C nucleotides, whereas *SCoVrC* genomes shows the highest percentages of C nucleotide. The highest percentages of A nucleotide were found for the two *PangSar* genomes, whereas the lowest percentage of A nucleotide was found for the RbBM48-31 genome. The lowest percentages of G nucleotide were detected for *SCoV2rC* and *PangSar* genomes.

**Figure 2.**
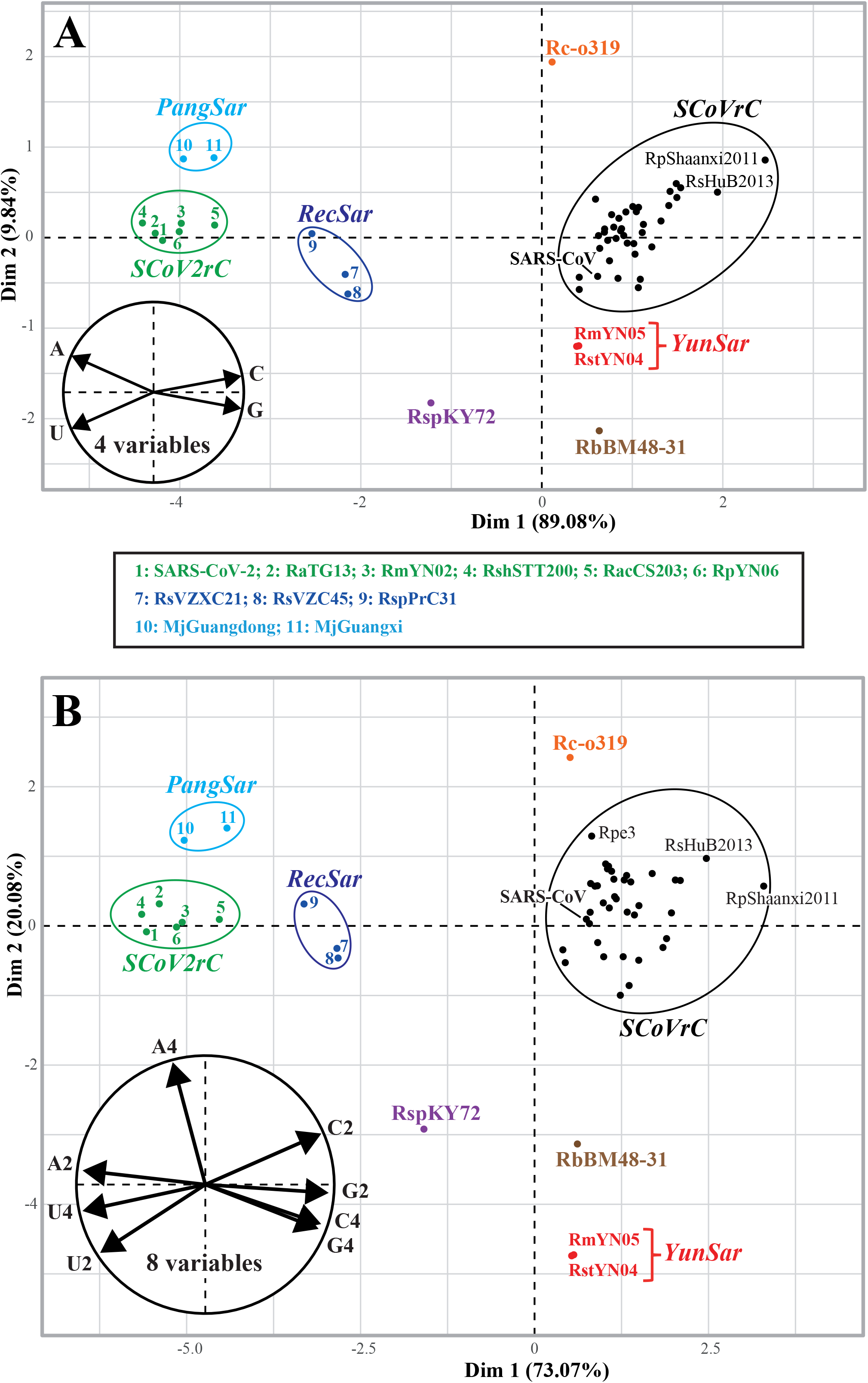
Variation in nucleotide composition at third codon-positions of *Sarbecovirus* genomes. The genomic alignment of 29,550 nt was used to calculate the frequency of the four bases (A, C, G and U) at third codon-positions (Table 2) and the 4 variables measured were then summarised by a principal component analysis (PCA; Figure **A**). The main graph represents the individual factor map based on 54 *Sarbecovirus* genomes. The eight groups of nucleotide composition are highlighted by different colours as defined in Figure 1. The small circular graph at the bottom left represents the variables factor map. The frequency of the four bases was also calculated either at four-fold degenerate third codon-positions or at two-fold degenerate third codon-positions for either purines (A *versus* G) or pyrimidines (C *versus* U) (Table 2). The PCA obtained using these 8 variables is shown in Figure **B**. The variables factor map is shown at the bottom left.

**Table 2.** Bases frequencies at third codon-positions (3CP)

| Third codon-positions | Bases | *SCoV2rC* | *PangSar* | *RecSar* | *SCoVrC* | *YunSar* | *Rc-o319* | *RbBM48-31* | *RspKY72* |
| --- | --- | --- | --- | --- | --- | --- | --- | --- | --- |
| All 3CP | A | 27.8-28.3 | 28.4-28.6 | 26.5-27.1 | 24.4-25.6 | 24.3 | 27.4 | 23.3 | 25.0 |
|  | C | 15.6-16.1 | 16.1-16.4 | 16.4-16.7 | 18.5-20.5 | 18.3 | 18.5 | 18.3 | 16.4 |
|  | G | 12.4-13.0 | 12.6-12.7 | 13.8-14.5 | 15.5-16.4 | 15.9 | 15.4 | 16.1 | 15.6 |
|  | U | 43.1-43.6 | 42.4-42.7 | 42.4-42.6 | 38.4-41.0 | 41.6 | 38.7 | 42.3 | 43.0 |
|  | A+U° | 70.9-71.9 | 70.8-71.3 | 69.1-69.5 | 63.1-65.8 | 65.8-65.9 | 66.1 | 65.6 | 68.0 |
| 4-fold degenerate 3CP | A | 28.9-29.3 | 29.9-30.0 | 28.5-29.3 | 27.5-29.8 | 24.3 | 31.1 | 25.1 | 26.3 |
|  | C | 13.5-14.2 | 13.6-13.8 | 14.2-14.5 | 16.1-17.8 | 19.2 | 16.1 | 17.1 | 15.3 |
|  | G | 6.4-6.9 | 6.2-6.7 | 7.2-8.0 | 8.1-10.2 | 9.6-9.7 | 8.6 | 9.1 | 9.2 |
|  | U | 49.8-50.9 | 49.4-50.3 | 49.0-49.2 | 44.1-46.9 | 46.8-46.9 | 44.2 | 48.8 | 49.2 |
|  | A+U° | 78.8-80.1 | 79.5-80.2 | 77.7-78.3 | 72.2-75.4 | 71.1-71.2 | 75.3 | 73.9 | 75.5 |
| 2-fold degenerate 3CP | A (vs G)* | 66.7-68.5 | 67.5-67.6 | 61.7-63.5 | 55.3-58.5 | 57.7-57.8 | 58.4 | 54.8 | 59.6 |
|  | C (vs U)* | 31.6-32.9 | 33.9-34.7 | 33.6-33.9 | 37.1-42.1 | 32.8 | 32.3 | 35.8 | 38.7 |

Highest percentages are highlighted in green; lowest percentages are highlighted in red.
°: variables not used in the PCAs of Figure 2; \*: the two variables were used in the PCA of Figure 2.

The nucleotide composition was also analysed at four-fold and two-fold degenerate third codon-positions (Table 2; CSV file available at https://osf.io/XXXX/). The eight variables were summarised by a PCA based on the first two PCs, which contribute 73.07% and 20.08% of the total variance, respectively (Figure 2B). The results confirmed the separation into eight groups, and some of them can be diagnosed by specific features, such as *YunSar* (less A nucleotides and more C nucleotides at four-fold degenerate third codon-positions), Rc-o319 (highest percentage of A nucleotide at four-fold degenerate third codon-positions), and RbBM48-31 (lowest percentage of A nucleotide at two-fold degenerate third codon-positions). The group uniting *SCoV2rC* and *PangSar* exhibits less G nucleotides and more U nucleotides at four-fold degenerate third codon-positions, as well as more A nucleotides at two-fold degenerate third codon-positions. For all variables at third codon-positions, *RecSar* genomes show intermediate values between *SCoV2rC* and *SCoVrC* genomes (Table 2).

### Dinucleotide composition

The dinucleotide frequencies at second and third codon-positions (P23) and at third and first codon-positions (P31) are provided in Table 3 (CSV files available at https://osf.io/XXXX/). For P23 and P31, the 16 variables were summarised by a PCA based on the first two PCs: for P23, they contribute 55.99% and 25.98% of the total variance, respectively (Figure 3A); for P31, they contribute 64.13% and 17.83% of the total variance, respectively (Figure 3B). The results showed a separation into the same eight groups previously identified, and some of them can be diagnosed by special features: *PangSar* genomes show less CG and GG at P23, and less CC and GU at P31; *SCoVrC* genomes are characterised by the highest precentages of GC and UC at P23; *SCoV2rC* genomes are characterised by the lowest precentages of GA at P31; *YunSar* genomes exhibit less AA, CA and GA at P23, less UC and UG at P31, more CC, CG, GU, and UA at P23, and more CG, GG, UA at P31; the Rc-o319 genome is the poorest in CU and UU at P23, whereas it is the richest in CA and GA at P23, and in AU at P31; the RbBM48-31 genome is the poorest in AA and AU at P31; the RspKY72 genome is the poorest in UC at P23 and in CU at P31, whereas it is the richest in AU and UG at P23, and in GU and UU at P31. The *SCoV2rC* and *PangSar* genomes are characterised by the highest percentages of AA at P23 and AC at P31, and by the lowest percentages of AG and CG at P23, and CC, CG, GA, GC, and GG at P31. The group uniting *SCoV2rC*, *PangSar* and *RecSar* show more AA at P31, less GC at P23, and less CC, CG, GC, and GG at P31. For all dinucleotides, *RecSar* genomes exhibit intermediate frequencies between *SCoV2rC* and *SCoVrC* genomes (Table 3).

**Figure 3.**
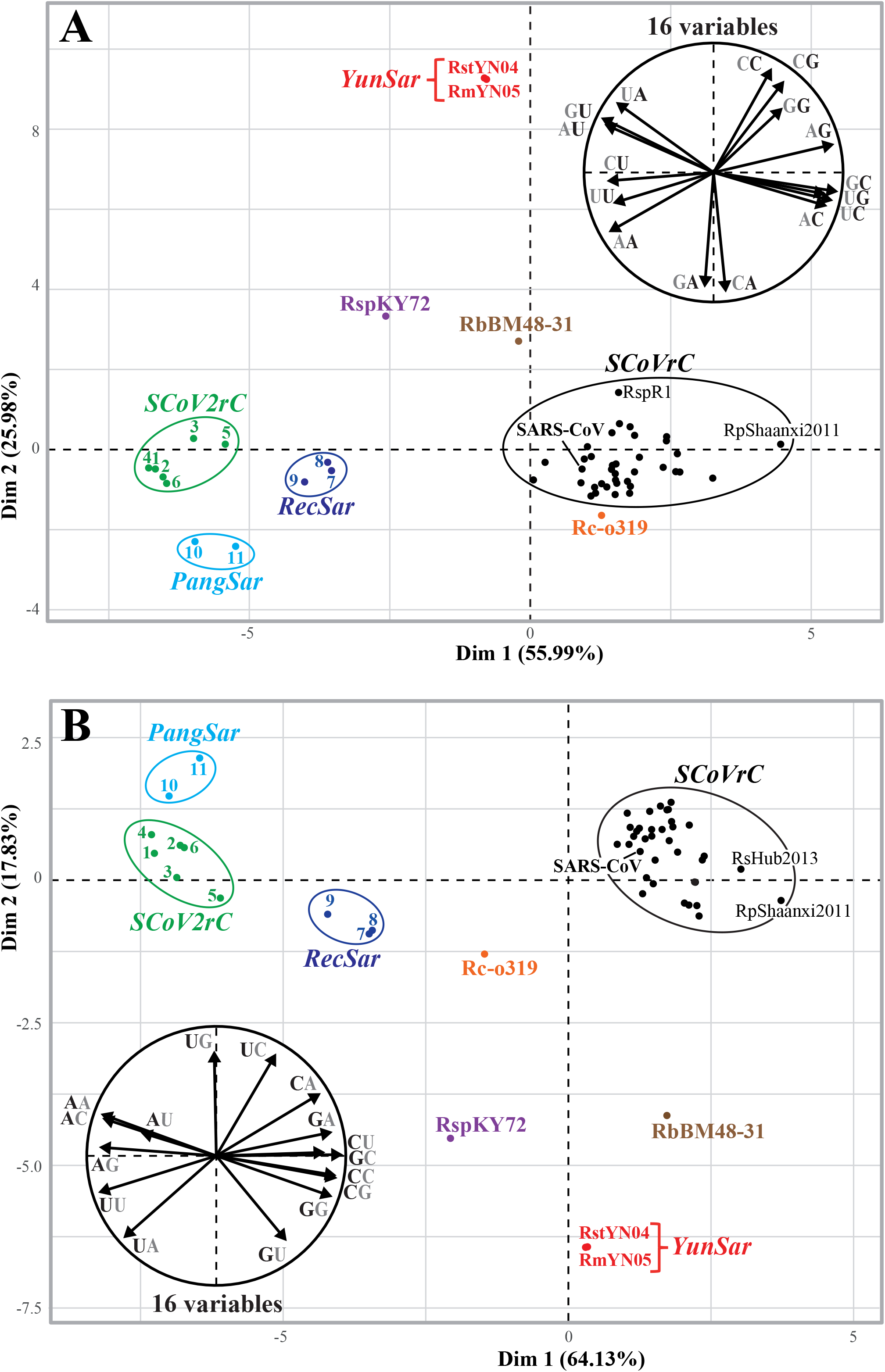
Variation in dinucleotide composition among *Sarbecovirus* genomes. The genomic alignment of 29,550 nt was used to calculate the frequencies of the 16 possible dinucleotides at second and third codon positions (P23) (Table 3) and the variables were summarised by a principal component analysis (PCA; Figure **A**). The main graph represents the individual factor map based on 54 *Sarbecovirus* genomes. The eight groups of dinucleotide composition are highlighted by different colours as defined in Figure 1. The small circular graph at the top right represents the variables factor map. The dinucleotide frequencies at third and first codon positions (P31) were also calculated (Table 3) and the 16 variables were summarised by a PCA (Figure **B**). The variables factor map is shown at the bottom left.

**Table 3.**
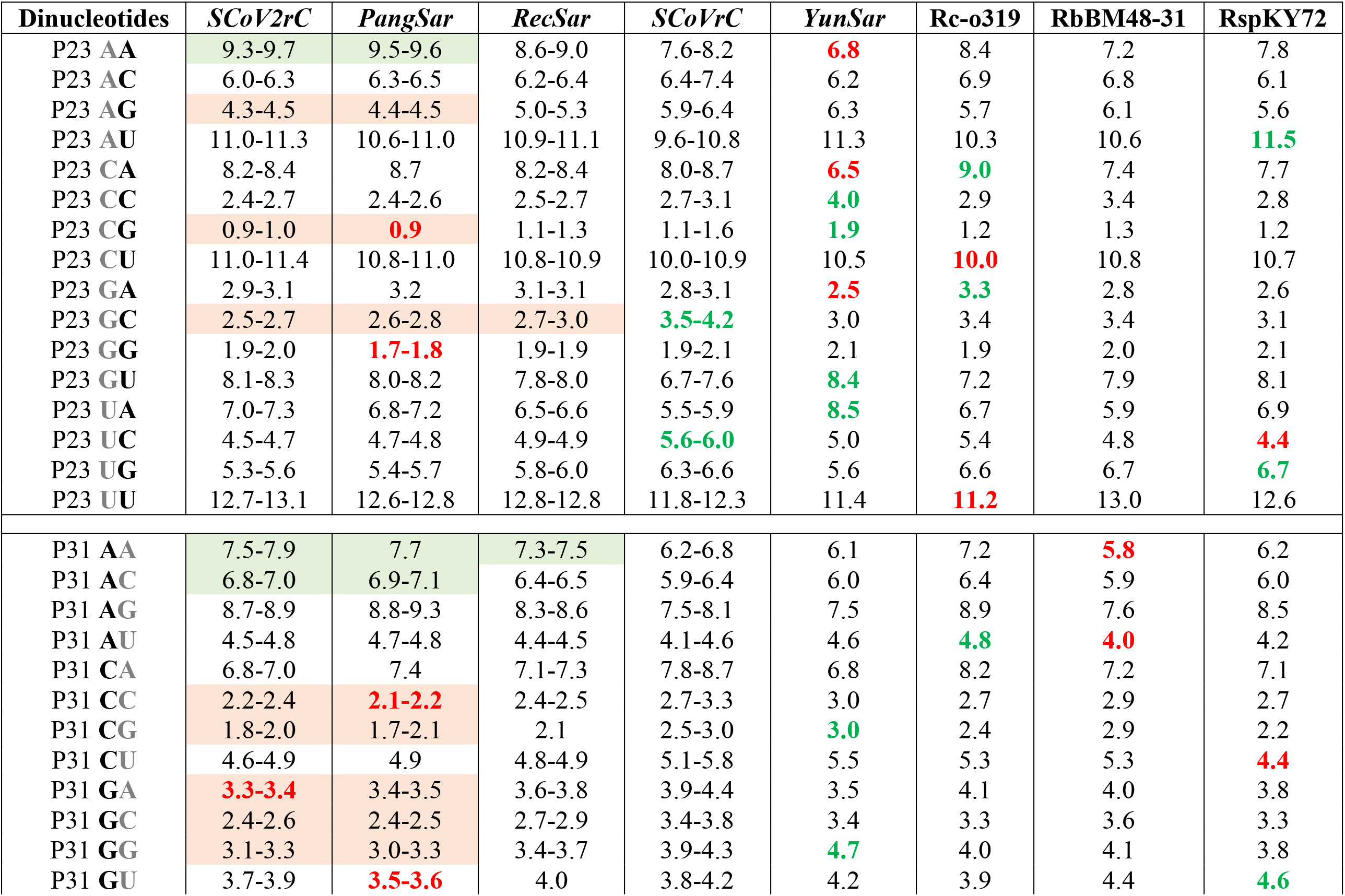
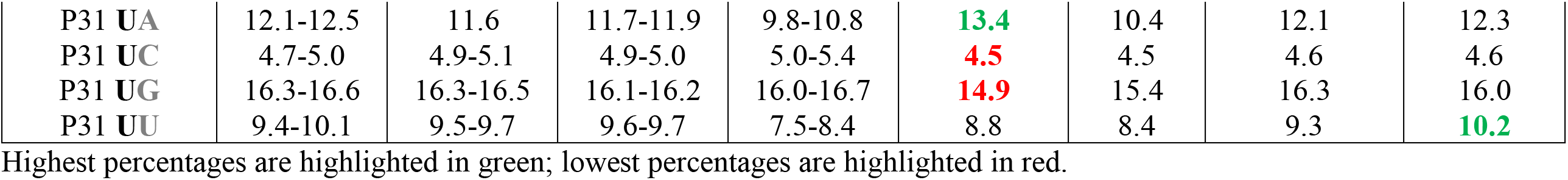
Dinucleotides frequencies at second and third codon-positions (P23) and at third and first codon-positions (P31)

| **Dinucleotides** | ***SCoV2rC*** | ***PangSar*** | ***RecSar*** | ***SCoVrC*** | ***YunSar*** | ***Rc-o319*** | ***RbBM48-31*** | ***RspKY72*** |
| --- | --- | --- | --- | --- | --- | --- | --- | --- |
| P23 **AA** | 9.3-9.7 | 9.5-9.6 | 8.6-9.0 | 7.6-8.2 | **6.8** | 8.4 | 7.2 | 7.8 |
| P23 **AC** | 6.0-6.3 | 6.3-6.5 | 6.2-6.4 | 6.4-7.4 | 6.2 | 6.9 | 6.8 | 6.1 |
| P23 **AG** | 4.3-4.5 | 4.4-4.5 | 5.0-5.3 | 5.9-6.4 | 6.3 | 5.7 | 6.1 | 5.6 |
| P23 **AU** | 11.0-11.3 | 10.6-11.0 | 10.9-11.1 | 9.6-10.8 | 11.3 | 10.3 | 10.6 | **11.5** |
| P23 **CA** | 8.2-8.4 | 8.7 | 8.2-8.4 | 8.0-8.7 | **6.5** | **9.0** | 7.4 | 7.7 |
| P23 **CC** | 2.4-2.7 | 2.4-2.6 | 2.5-2.7 | 2.7-3.1 | **4.0** | 2.9 | 3.4 | 2.8 |
| P23 **CG** | 0.9-1.0 | **0.9** | 1.1-1.3 | 1.1-1.6 | **1.9** | 1.2 | 1.3 | 1.2 |
| P23 **CU** | 11.0-11.4 | 10.8-11.0 | 10.8-10.9 | 10.0-10.9 | 10.5 | **10.0** | 10.8 | 10.7 |
| P23 **GA** | 2.9-3.1 | 3.2 | 3.1-3.1 | 2.8-3.1 | **2.5** | **3.3** | 2.8 | 2.6 |
| P23 **GC** | 2.5-2.7 | 2.6-2.8 | 2.7-3.0 | **3.5-4.2** | 3.0 | 3.4 | 3.4 | 3.1 |
| P23 **GG** | 1.9-2.0 | **1.7-1.8** | 1.9-1.9 | 1.9-2.1 | 2.1 | 1.9 | 2.0 | 2.1 |
| P23 **GU** | 8.1-8.3 | 8.0-8.2 | 7.8-8.0 | 6.7-7.6 | **8.4** | 7.2 | 7.9 | 8.1 |
| P23 **UA** | 7.0-7.3 | 6.8-7.2 | 6.5-6.6 | 5.5-5.9 | **8.5** | 6.7 | 5.9 | 6.9 |
| P23 **UC** | 4.5-4.7 | 4.7-4.8 | 4.9-4.9 | **5.6-6.0** | 5.0 | 5.4 | 4.8 | **4.4** |
| P23 **UG** | 5.3-5.6 | 5.4-5.7 | 5.8-6.0 | 6.3-6.6 | 5.6 | 6.6 | 6.7 | **6.7** |
| P23 **UU** | 12.7-13.1 | 12.6-12.8 | 12.8-12.8 | 11.8-12.3 | 11.4 | **11.2** | 13.0 | 12.6 |
| P31 **AA** | 7.5-7.9 | 7.7 | 7.3-7.5 | 6.2-6.8 | 6.1 | 7.2 | **5.8** | 6.2 |
| P31 **AC** | 6.8-7.0 | 6.9-7.1 | 6.4-6.5 | 5.9-6.4 | 6.0 | 6.4 | 5.9 | 6.0 |
| P31 **AG** | 8.7-8.9 | 8.8-9.3 | 8.3-8.6 | 7.5-8.1 | 7.5 | 8.9 | 7.6 | 8.5 |
| P31 **AU** | 4.5-4.8 | 4.7-4.8 | 4.4-4.5 | 4.1-4.6 | 4.6 | **4.8** | **4.0** | 4.2 |
| P31 **CA** | 6.8-7.0 | 7.4 | 7.1-7.3 | 7.8-8.7 | 6.8 | 8.2 | 7.2 | 7.1 |
| P31 **CC** | 2.2-2.4 | **2.1-2.2** | 2.4-2.5 | 2.7-3.3 | 3.0 | 2.7 | 2.9 | 2.7 |
| P31 **CG** | 1.8-2.0 | 1.7-2.1 | 2.1 | 2.5-3.0 | **3.0** | 2.4 | 2.9 | 2.2 |
| P31 **CU** | 4.6-4.9 | 4.9 | 4.8-4.9 | 5.1-5.8 | 5.5 | 5.3 | 5.3 | **4.4** |
| P31 **GA** | **3.3-3.4** | 3.4-3.5 | 3.6-3.8 | 3.9-4.4 | 3.5 | 4.1 | 4.0 | 3.8 |
| P31 **GC** | 2.4-2.6 | 2.4-2.5 | 2.7-2.9 | 3.4-3.8 | 3.4 | 3.3 | 3.6 | 3.3 |
| P31 **GG** | 3.1-3.3 | 3.0-3.3 | 3.4-3.7 | 3.9-4.3 | **4.7** | 4.0 | 4.1 | 3.8 |
| P31 **GU** | 3.7-3.9 | **3.5-3.6** | 4.0 | 3.8-4.2 | 4.2 | 3.9 | 4.4 | **4.6** |
| P31 **UA** | 12.1-12.5 | 11.6 | 11.7-11.9 | 9.8-10.8 | **13.4** | 10.4 | 12.1 | 12.3 |
| P31 **UC** | 4.7-5.0 | 4.9-5.1 | 4.9-5.0 | 5.0-5.4 | **4.5** | 4.5 | 4.6 | 4.6 |
| P31 **UG** | 16.3-16.6 | 16.3-16.5 | 16.1-16.2 | 16.0-16.7 | **14.9** | 15.4 | 16.3 | 16.0 |
| P31 **UU** | 9.4-10.1 | 9.5-9.7 | 9.6-9.7 | 7.5-8.4 | 8.8 | 8.4 | 9.3 | **10.2** |

Highest percentages are highlighted in green; lowest percentages are highlighted in red.

### Codon usage

The 59 variables corresponding to the relative frequencies between synonymous codons of 18 different amino acids (all except M and W) were summarised in Table 4 (CSV file available at https://osf.io/XXXX/). The first two dimensions of the PCA contribute 44.93% and 19.37% of the total variance, respectively (Figure 4). Here again the results confirmed the division into the eigth groups previsouly identified, and some of them can be diagnosed by special features in codon usage: *PangSar* genomes have the lowest percentages for GCC Alanine codon; *SCoVrC* genomes have the lowest percentages for AUA Isoleucine codon, UUA Leucine codon and GUU Glycine codon, and the highest percentages for AUC Isoleucine codon, CUC and CUG Leucine codons; *YunSar* genomes have an atypical codon composition, as they show the lowest percentages for codons AAA, ACA, AUU, CUU, GCA, GGA, UAC, UCA, and UUG and the highest percentages for codons ACC, ACG, AUA, CCC, GCC, UCC, UCG, and UUA; the Rc-o319 genome is the poorest in ACU Threonine codons and GUU Valine codons, whereas it is the richest in ACA Threonine codons, GCA Alanine codons, and GUA Valine codons; the RbBM48-31 genome exhibits the lowest percentages for CAA Glutamine codon, CCA Proline codon, and GUA Valine codon; and the RspKY72 genome shows the lowest percentages for AUC Isoleucine codon, CAC Histidine codon, CGA Arginine codon, and CUC Leucine codon, and the highest percentages for CCU Proline codon, AGG and CGG Arginine codons, and UUG Leucine codon. The *SCoV2rC* and *PangSar* genomes are characterised by the lowest percentages for CCC Proline codon and GGC Glycine codon, and by the highest percentages for AAA Lysine codon, CAA Glutamine codon, and GAA Glutamate codon. The group uniting *SCoV2rC*, *PangSar* and *RecSar* is characterised by the lowest percantages for GUG Valine codons. For all variables, *RecSar* genomes show intermediate values between *SCoV2rC* and *SCoVrC* genomes (Table 4).

**Figure 4.**
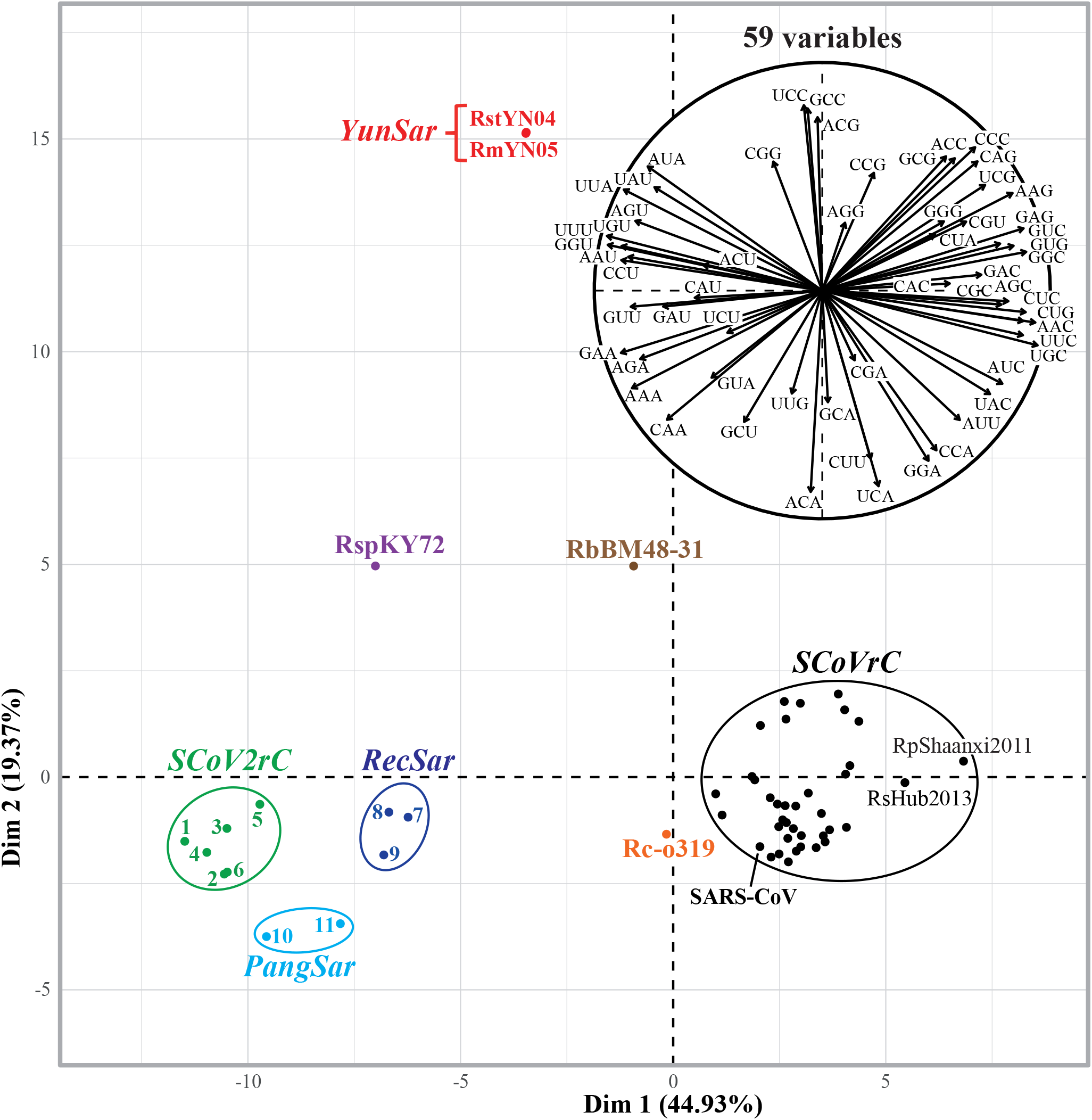
Variation in codon usage among *Sarbecovirus* genomes. The alignment of 29,550 nt was used to calculate the frequencies of synonymous codons for all amino acids except M and W that are encoded by a single codon (Table 4). The 59 variables were then summarised by a principal component analysis (PCA). The main graph represents the individual factor map based on 54 *Sarbecovirus* genomes. The eight groups of codon usage are highlighted by different colours as defined in Figure 1. The circular graph at the top right represents the variables factor map.

**Table 4.**
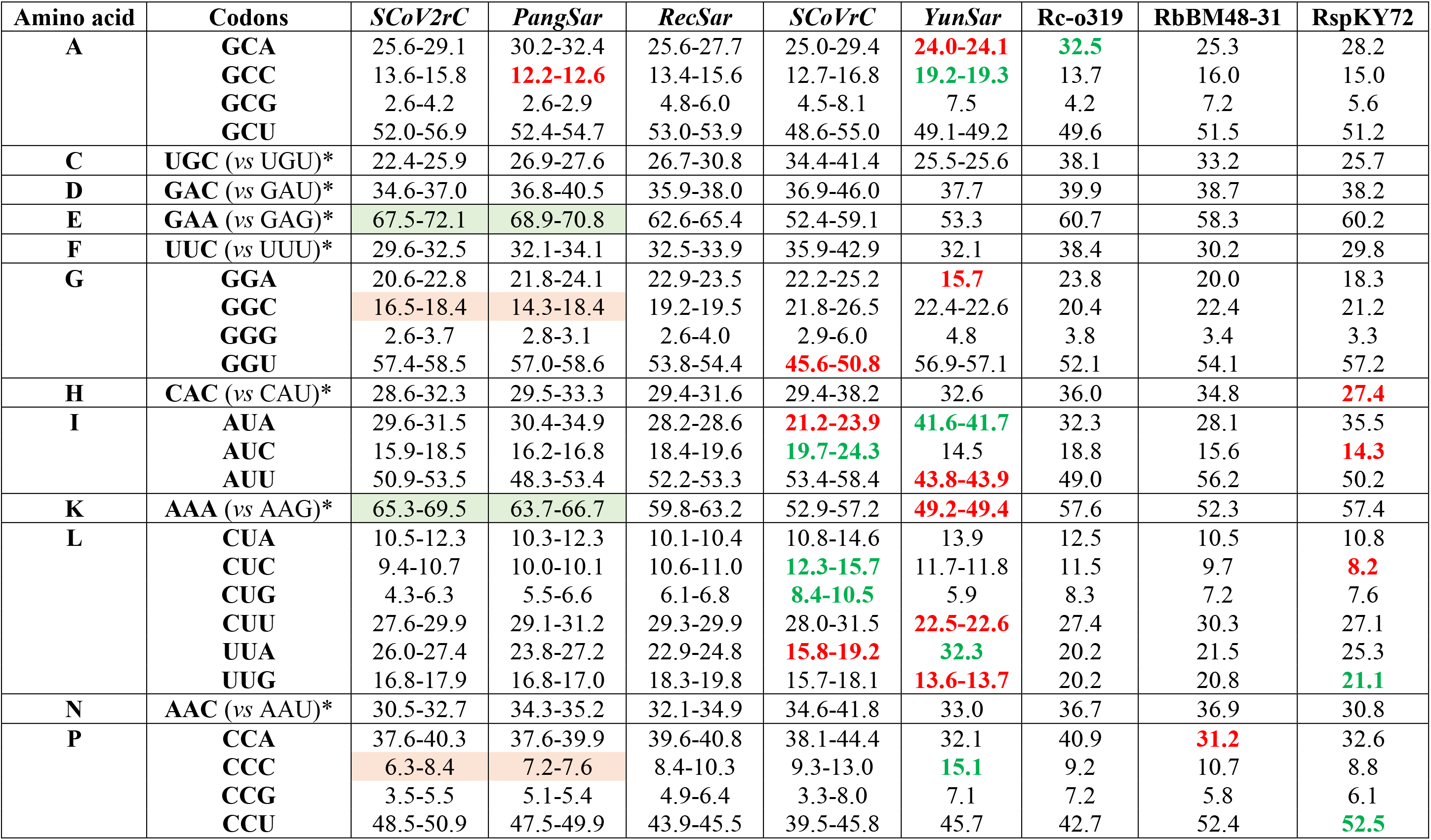
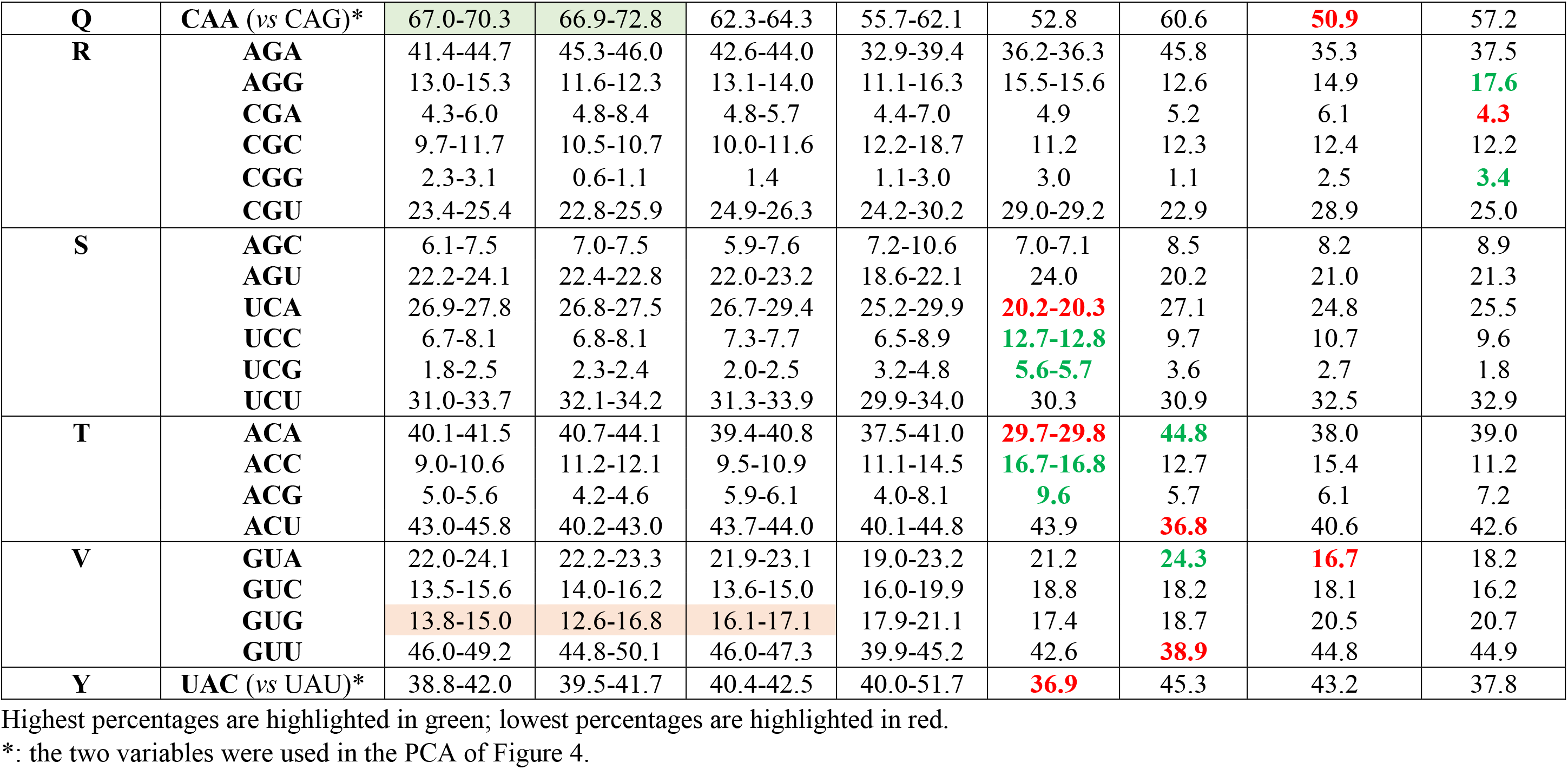
**Relative codon frequencies for all amino acids** (except M and W that are encoded by a single codon)

| Amino acid | Codons | *SCoV2rC* | *PangSar* | *RecSar* | *SCoVrC* | *YunSar* | Rc-o319 | RbBM48-31 | RspKY72 |
| --- | --- | --- | --- | --- | --- | --- | --- | --- | --- |
| **A** | **GCA** | 25.6-29.1 | 30.2-32.4 | 25.6-27.7 | 25.0-29.4 | **24.0-24.1** | **32.5** | 25.3 | 28.2 |
|  | **GCC** | 13.6-15.8 | **12.2-12.6** | 13.4-15.6 | 12.7-16.8 | **19.2-19.3** | 13.7 | 16.0 | 15.0 |
|  | **GCG** | 2.6-4.2 | 2.6-2.9 | 4.8-6.0 | 4.5-8.1 | 7.5 | 4.2 | 7.2 | 5.6 |
|  | **GCU** | 52.0-56.9 | 52.4-54.7 | 53.0-53.9 | 48.6-55.0 | 49.1-49.2 | 49.6 | 51.5 | 51.2 |
| **C** | **UGC** (vs UGU)* | 22.4-25.9 | 26.9-27.6 | 26.7-30.8 | 34.4-41.4 | 25.5-25.6 | 38.1 | 33.2 | 25.7 |
| **D** | **GAC** (vs GAU)* | 34.6-37.0 | 36.8-40.5 | 35.9-38.0 | 36.9-46.0 | 37.7 | 39.9 | 38.7 | 38.2 |
| **E** | **GAA** (vs GAG)* | 67.5-72.1 | 68.9-70.8 | 62.6-65.4 | 52.4-59.1 | 53.3 | 60.7 | 58.3 | 60.2 |
| **F** | **UUC** (vs UUU)* | 29.6-32.5 | 32.1-34.1 | 32.5-33.9 | 35.9-42.9 | 32.1 | 38.4 | 30.2 | 29.8 |
| **G** | **GGA** | 20.6-22.8 | 21.8-24.1 | 22.9-23.5 | 22.2-25.2 | **15.7** | 23.8 | 20.0 | 18.3 |
|  | **GGC** | 16.5-18.4 | 14.3-18.4 | 19.2-19.5 | 21.8-26.5 | 22.4-22.6 | 20.4 | 22.4 | 21.2 |
|  | **GGG** | 2.6-3.7 | 2.8-3.1 | 2.6-4.0 | 2.9-6.0 | 4.8 | 3.8 | 3.4 | 3.3 |
|  | **GGU** | 57.4-58.5 | 57.0-58.6 | 53.8-54.4 | **45.6-50.8** | 56.9-57.1 | 52.1 | 54.1 | 57.2 |
| **H** | **CAC** (vs CAU)* | 28.6-32.3 | 29.5-33.3 | 29.4-31.6 | 29.4-38.2 | 32.6 | 36.0 | 34.8 | **27.4** |
| **I** | **AUA** | 29.6-31.5 | 30.4-34.9 | 28.2-28.6 | **21.2-23.9** | **41.6-41.7** | 32.3 | 28.1 | 35.5 |
|  | **AUC** | 15.9-18.5 | 16.2-16.8 | 18.4-19.6 | **19.7-24.3** | 14.5 | 18.8 | 15.6 | **14.3** |
|  | **AUU** | 50.9-53.5 | 48.3-53.4 | 52.2-53.3 | 53.4-58.4 | **43.8-43.9** | 49.0 | 56.2 | 50.2 |
| **K** | **AAA** (vs AAG)* | 65.3-69.5 | 63.7-66.7 | 59.8-63.2 | 52.9-57.2 | **49.2-49.4** | 57.6 | 52.3 | 57.4 |
| **L** | **CUA** | 10.5-12.3 | 10.3-12.3 | 10.1-10.4 | 10.8-14.6 | 13.9 | 12.5 | 10.5 | 10.8 |
|  | **CUC** | 9.4-10.7 | 10.0-10.1 | 10.6-11.0 | **12.3-15.7** | 11.7-11.8 | 11.5 | 9.7 | **8.2** |
|  | **CUG** | 4.3-6.3 | 5.5-6.6 | 6.1-6.8 | **8.4-10.5** | 5.9 | 8.3 | 7.2 | 7.6 |
|  | **CUU** | 27.6-29.9 | 29.1-31.2 | 29.3-29.9 | 28.0-31.5 | **22.5-22.6** | 27.4 | 30.3 | 27.1 |
|  | **UUA** | 26.0-27.4 | 23.8-27.2 | 22.9-24.8 | **15.8-19.2** | **32.3** | 20.2 | 21.5 | 25.3 |
|  | **UUG** | 16.8-17.9 | 16.8-17.0 | 18.3-19.8 | 15.7-18.1 | **13.6-13.7** | 20.2 | 20.8 | **21.1** |
| **N** | **AAC** (vs AAU)* | 30.5-32.7 | 34.3-35.2 | 32.1-34.9 | 34.6-41.8 | 33.0 | 36.7 | 36.9 | 30.8 |
| **P** | **CCA** | 37.6-40.3 | 37.6-39.9 | 39.6-40.8 | 38.1-44.4 | 32.1 | 40.9 | **31.2** | 32.6 |
|  | **CCC** | 6.3-8.4 | 7.2-7.6 | 8.4-10.3 | 9.3-13.0 | **15.1** | 9.2 | 10.7 | 8.8 |
|  | **CCG** | 3.5-5.5 | 5.1-5.4 | 4.9-6.4 | 3.3-8.0 | 7.1 | 7.2 | 5.8 | 6.1 |
|  | **CCU** | 48.5-50.9 | 47.5-49.9 | 43.9-45.5 | 39.5-45.8 | 45.7 | 42.7 | 52.4 | **52.5** |
| **Q** | **CAA (vs CAG)\*** | 67.0-70.3 | 66.9-72.8 | 62.3-64.3 | 55.7-62.1 | 52.8 | 60.6 | **50.9** | 57.2 |
| **R** | **AGA** | 41.4-44.7 | 45.3-46.0 | 42.6-44.0 | 32.9-39.4 | 36.2-36.3 | 45.8 | 35.3 | 37.5 |
|  | **AGG** | 13.0-15.3 | 11.6-12.3 | 13.1-14.0 | 11.1-16.3 | 15.5-15.6 | 12.6 | 14.9 | **17.6** |
|  | **CGA** | 4.3-6.0 | 4.8-8.4 | 4.8-5.7 | 4.4-7.0 | 4.9 | 5.2 | 6.1 | **4.3** |
|  | **CGC** | 9.7-11.7 | 10.5-10.7 | 10.0-11.6 | 12.2-18.7 | 11.2 | 12.3 | 12.4 | 12.2 |
|  | **CGG** | 2.3-3.1 | 0.6-1.1 | 1.4 | 1.1-3.0 | 3.0 | 1.1 | 2.5 | **3.4** |
|  | **CGU** | 23.4-25.4 | 22.8-25.9 | 24.9-26.3 | 24.2-30.2 | 29.0-29.2 | 22.9 | 28.9 | 25.0 |
| **S** | **AGC** | 6.1-7.5 | 7.0-7.5 | 5.9-7.6 | 7.2-10.6 | 7.0-7.1 | 8.5 | 8.2 | 8.9 |
|  | **AGU** | 22.2-24.1 | 22.4-22.8 | 22.0-23.2 | 18.6-22.1 | 24.0 | 20.2 | 21.0 | 21.3 |
|  | **UCA** | 26.9-27.8 | 26.8-27.5 | 26.7-29.4 | 25.2-29.9 | **20.2-20.3** | 27.1 | 24.8 | 25.5 |
|  | **UCC** | 6.7-8.1 | 6.8-8.1 | 7.3-7.7 | 6.5-8.9 | **12.7-12.8** | 9.7 | 10.7 | 9.6 |
|  | **UCG** | 1.8-2.5 | 2.3-2.4 | 2.0-2.5 | 3.2-4.8 | **5.6-5.7** | 3.6 | 2.7 | 1.8 |
|  | **UCU** | 31.0-33.7 | 32.1-34.2 | 31.3-33.9 | 29.9-34.0 | 30.3 | 30.9 | 32.5 | 32.9 |
| **T** | **ACA** | 40.1-41.5 | 40.7-44.1 | 39.4-40.8 | 37.5-41.0 | **29.7-29.8** | **44.8** | 38.0 | 39.0 |
|  | **ACC** | 9.0-10.6 | 11.2-12.1 | 9.5-10.9 | 11.1-14.5 | **16.7-16.8** | 12.7 | 15.4 | 11.2 |
|  | **ACG** | 5.0-5.6 | 4.2-4.6 | 5.9-6.1 | 4.0-8.1 | **9.6** | 5.7 | 6.1 | 7.2 |
|  | **ACU** | 43.0-45.8 | 40.2-43.0 | 43.7-44.0 | 40.1-44.8 | 43.9 | **36.8** | 40.6 | 42.6 |
| **V** | **GUA** | 22.0-24.1 | 22.2-23.3 | 21.9-23.1 | 19.0-23.2 | 21.2 | **24.3** | **16.7** | 18.2 |
|  | **GUC** | 13.5-15.6 | 14.0-16.2 | 13.6-15.0 | 16.0-19.9 | 18.8 | 18.2 | 18.1 | 16.2 |
|  | **GUG** | 13.8-15.0 | 12.6-16.8 | 16.1-17.1 | 17.9-21.1 | 17.4 | 18.7 | 20.5 | 20.7 |
|  | **GUU** | 46.0-49.2 | 44.8-50.1 | 46.0-47.3 | 39.9-45.2 | 42.6 | **38.9** | 44.8 | 44.9 |
| **Y** | **UAC (vs UAU)\*** | 38.8-42.0 | 39.5-41.7 | 40.4-42.5 | 40.0-51.7 | **36.9** | 45.3 | 43.2 | 37.8 |

Highest percentages are highlighted in green; lowest percentages are highlighted in red.
\*: the two variables were used in the PCA of Figure 4.

## Discussion

### Phylogenetic evidence for past events of recombination between divergent *Sarbecovirus* genomes

The phylogeny of sarbecoviruses is difficult to apprehend because their evolutionnary history has been punctated by multiple events of genomic recombination (Boni et al. 2020). As a consequence, different genomic regions can bring incongruent phylogenetic signals, and different methods of tree reconstruction can provide discordant tree topologies. For that reason, the topology of the bootstrap consensus tree (Figure 1A) differs from the best ML tree (data not shown; NEWICK tree available at https://osf.io/XXXX/) while both trees were constructed from the same whole genomic alignment. In addition, two gene trees can show high support for distinct evolutionary histories. For instance, the *RecSar* group was found to be monophyletic (BP = 100) and closely related to RpYN06 (bat *SCoV2rC*) (BP = 100) in the Spike gene tree (Figure 1D), whereas *RecSar* appeared as polyphyletic in the RdRp gene tree, RsZC45 and RsZXC21 (both from Zhejiang) being closely related to four *SCoVrC* viruses (RmoLongquan 140, RsHKU3-1, -7, and -12) (BP = 100) and RspPrC31 (from Yunnan) being enclosed into a large clade of *SCoVrCs* containing SARS-CoV and several bat viruses (BP = 89). In agreement with previous studies (Hu et al. 2018; Boni et al. 2020; Li et al. 2021), these results indicate that the three *RecSar* viruses (RspPrC31, RsZC45, and RsZXC21) have emerged through recombination between *SCoVrC* and *SCoV2rC* genomes. The genomic recombination between these two divergent *Sarbecovirus* lineages may have occurred preferrentially in the geographic area including southern Yunnan, northern Laos and northern Vietnam, where the ecological niches of *SCoVrC* and *SCoV2rC* slightly overlap (Hassanin et al. 2021b). It can be also proposed that divergent or recombined variants of *Sarbecovirus* may have dispersed from Yunnan to eastern China through several generations of bats via occasional contacts in caves between populations usually found in different regions. Such a scenario involving a diffusion over several decades of new *Sarbecovirus* variants from Yunnan to other provinces of China should be tested with additional data.

There is also some phylogenetic evidence of past events of genomic recombination involving the ancestors of MjGuangdong, *SCoV2rC* and *YunSar*: in the RdRp gene tree (Figure 1C), *SCoV2rC* is monophyletic (BP = 84), whereas *YunSar* appears as the sister-group of MjGuangdong (BP = 99); in the Spike gene tree, *YunSar* is divergent from *SCoVrC*, *SCoV2rC*, *PangSar*, *RecSar*, Rc-o319 (BP = 100; Figure 1D), and MjGuangdong is related to RshSTT200, RaTG13, SARS-CoV-2 and MjGuangxi (BP = 100). Here again, these past events of genomic recombination between viruses from different lineages may have occurred in Yunnan, as *YunSar* and RaTG13 were collected in this province (Zhou H. et al. 2020, 2021), or alternatively in Southeast Asia, as RshSTT200 was described from northern Cambodia (Hul et al. 2021) and MjGuangdong and MjGuangxi came from Sunda pangolins collected in unknown localities of Southeast Asia (Hassanin et al. 2021a).

The data currently available for *Sarbecovirus* genomes indicate that the Yunnan province constitutes a key area for their evolution, as at least three groups (*SCoVrC*, *SCoV2rC* and *YunSar*) have been detected in this region of southern China (Han et al. 2019; Zhou H. et al. 2020, 2021; Zhou P. et al. 2020). This province is therefore the main place where recombination can occurr between divergent *Sarbecovirus* genomes. The three viral groups were all found in both *Rhinolophus affinis* and *Rhinolophus malayanus*, whereas two of them were detected in *Rhinolophus pusillus* (*SCoVrC* and *SCoV2rC*) and *Rhinolophus stheno* (*SCoVrC* and *YunSar*) (Figure 5). Interestingly, these four horseshoe bat species are distributed not only in Yunnan, but also in several Southeast Asian countries, such as Cambodia, Laos, Myanmar, Thailand, and Vietnam, where only two field investigations were carried out for searching bat coronaviruses: one in eastern Thailand in 2020 (Wacharapluesadee et al. 2021) and another in northern Cambodia in 2010 (Hul et al. 2021). Obviously, there is no doubt that future studies will describe more *Sarbecovirus* diversity in Southeast Asia.

**Figure 5.**
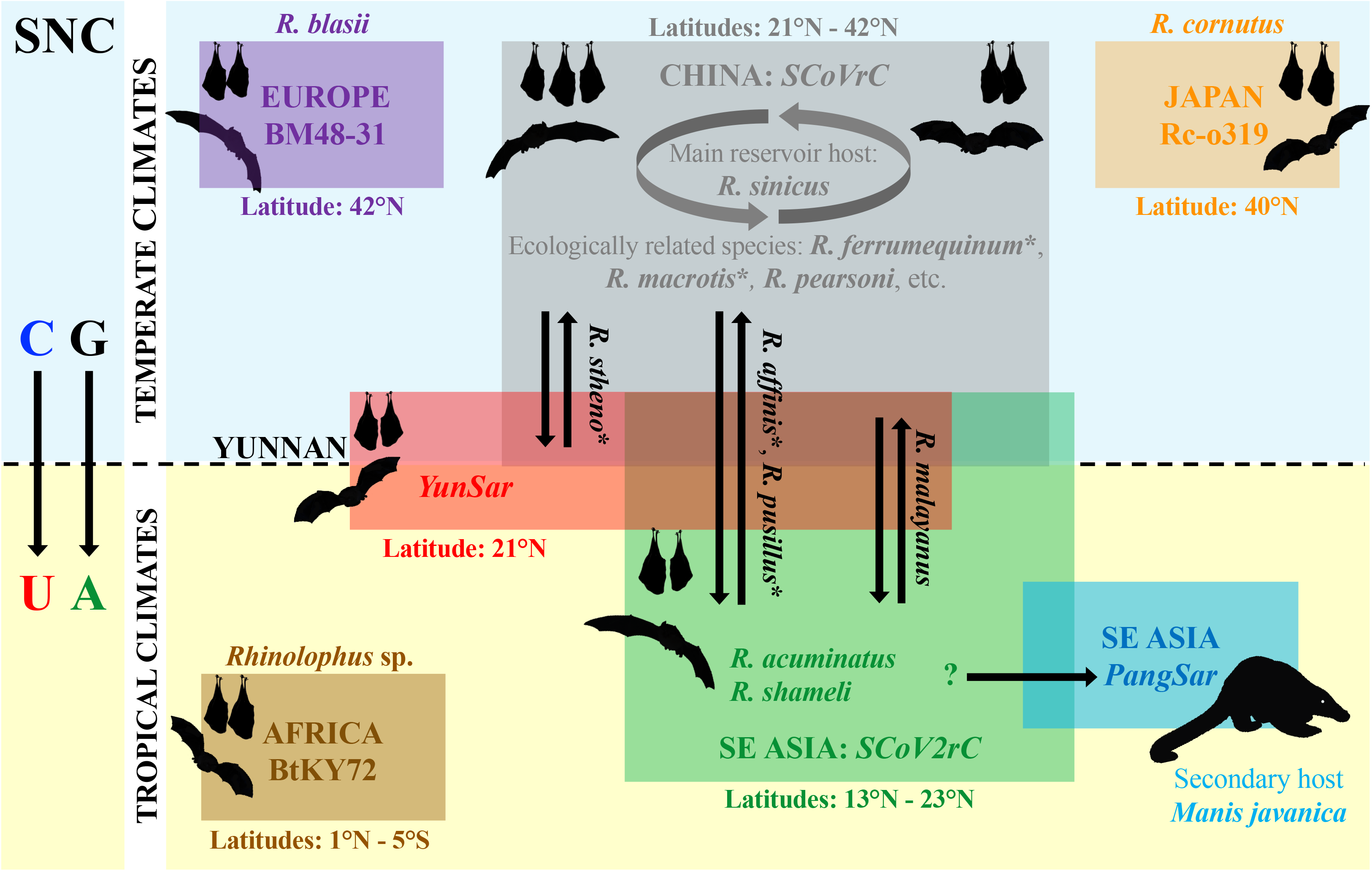
Host species and latitudinal distribution of the seven groups of *Sarbecovirus* genomes showing different synonymous nucleotide compositions. The seven groups of *Sarbecovirus* genomes are highlighted by coloured rectangles. The abbreviation “*R.*” is used for *Rhinolophus* species. The double arrows indicate *Rhinolophus* species from which several groups of *Sarbecovirus* were sequenced in previous studies. All species names concerned by taxonomic issues (see Table 1 for details) are followed by an asterisk. As shown in Table 2, the genomic bias in favour of A+U nucleotides is more marked for the groups of sarbecoviruses circulating in bats (or pangolins) from tropical latitudes (BtKY72 in Sub-Saharan Africa and *SCoV2rC* and *PangSar* in Southeast Asia) than for those from temperate latitudes (BM48-31 in Europe, *SCoVrC* in China, and Rc-o319 in Japan).

### Evidence for seven groups of SNC among *Sarbecovirus* genomes

In this study, all analyses of nucleotide composition, dinucleotide composition and codon usage (Figures 2-4) showed evidence for eight groups of *Sarbecovirus* genomes: (i) *SCoVrC*, including SARS-CoV and a large diversity of bat viruses from China; (ii) *SCoV2rC*, including SARS-CoV-2 and five bat viruses from Cambodia, Thailand and Yunnan, (iii) *PangSar*, which is composed of the two viruses detected in Sunda pangolins seized in the Chinese provinces of Guangdong (in 2019) and Guangxi (in 2017-2018); (iv) *RecSar*, which contains the three bat viruses showing evidence of past recombination between *SCoV2rC* and *SCoVrC* genomes; (v) *YunSar*, i.e., the two highly divergent bat viruses recently described from Yunnan by Zhou et al. (2021; RmYN05 and RstYN04); (vi) RbBM48-31, the bat virus from Bulgaria; (vii) RspKY72, the bat virus from Kenya; and (viii) Rc-o319, the bat virus from Japan. All these groups can be diagnosed by specific features (i.e., highest or lowest percentages) in nucleotide composition, dinucleotide composition, and/or codon usage (Tables 2-4). The only exception is *RecSar* for which all variables show intermediate values between those found for *SCoVrC* and *SCoV2rC*. Obviously, this is due to their origin by recombination between *SCoVrC* and *SCoV2rC* genomes (Boni et al. 2020; Li et al. 2021; present study).

The SNC is the primary factor explaining the results also observed at synonymous sites of dinucleotides and codons. Indeed, the two variables factor maps obtained from PCAs based on the nucleotide composition at third codon-positions (Figures 2A and 2B) revealed that the variance is mainly explained by the first dimension (89.08% and 73.07%, respectively), which separates *Sarbecorvirus* genomes showing the highest A+U content at third codon positions (at the left; *SCoV2rC*, *PangSar*, *RecSar*, and RspKY72; A+U (3CP) = 71.9-68.0%) from the other ones (at the right; *SCoVrC*, RbBM48-3, Rc-o319, and *YunSar*; A+U (3CP) = 66.1-63.1%). Similar results were found in the PCAs based on dinucleotide composition (Figures 3A and 3B) and codon usage (Figure 4), with *SCoV2rC* and *PangSar* genomes exhibiting a more marked bias towards A and U nucleotides (or against C+G nucleotides) at synonymous sites. All these results support a stronger mutational bias in *SCoV2rC* and *PangSar* genomes characterised by higher rates for C=>U and G=>A transitions than for the reverse mutations (U=>C and A=>G, respectively). Previous studies examining the nucleotide composition in SARS-CoV-2 genomes have all concluded to an over-representation of C=>U transitions (Rice et al. 2020; Matyášek et al. 2021). Several mechanisms have been proposed to account for the cytosine deficiency in the genome of sarbecoviruses, such as cytosine deamination resulting from the action of the host APOBEC system (Milewska et al. 2018), methylation of CpG dinucleotides (Xia 2020), or the limited availability of cytidine triphosphate (CTP), which is used not only for the viral RNA genome synthesis but also for the synthesis of the virus envelope, as well as translation and glycosylation of viral proteins (Ou et al. 2021). The bias in favor of C=>U transitions (over U=>C transitions) has been observed in a wide range of mammalian RNA viruses (Simmonds and Ansari 2021), indicating that it is the result of an asymmetrical mechanism shared by all RNA viruses infecting mammals. I suggest that the replication of viral RNA genomes, which is an asymmetrical process dependent of the pool of free nucleotides available in infected cells, can explain the seven SNC patterns here observed among *Sarbecovirus* genomes. From this point of view, it is important to note that the physiological concentrations of nucleotides were found to be highly variable among mammalian species and tissues, with the following means and standard deviations (in μM) published in Traut (1994): ATP = 3,152 ± 1,698 > UTP = 567 ± 460 > GTP = 468 ± 224 > CTP = 278 ± 242. When a mammalian cell divides, the synthesis of nucleotides is regulated at multiple levels to maintain constant concentrations of free nucleotides (Lane and Fan 2015). By contrast, the replication of viral RNA genomes always takes place in mammalian cells containing imbalanced nucleotide pools (A >> U > G > C) (Traut 1994), which is supposed to promote higher mutation rates for G=>A transitions (*versus* A=>G transitions) and to a lesser extent C=>U transitions (*versus* U=>C transitions). In addition, the availability of CTP is much more reduced when RNA viruses multiply in their host cells (Ou et al. 2021), increasing therefore the rate of C=>U transitions during viral replication. All these elements suggest that taxonomic variation in the concentrations of free nucleotides available in infected cells during viral RNA replication is the main evolutionary force behind the variation in SNC here observed among *Sarbecovirus* genomes.

From the taxonomic point of view, it is relevant to remark that all the seven SNC groups of *Sarbecovirus* genomes, except *PangSar* (but see next paragraph), were found in different species or species assemblages of horseshoe bats (family Rhinolophidae, genus *Rhinolophus*) (Figure 5). The genus *Rhinolophus* currently includes between 92 and 109 insectivorous bat species (Burgin et al. 2020; IUCN 2021) that inhabit temperate and tropical regions of the Old World, with a higher biodiversity in Asia (63-68 out of the 92-109 described species) than in Africa (34-38 species), Europe (5 species) and Oceania (5 species). All *Rhinolophus* species in which sarbecoviruses were detected in previous studies (Table 1) are cave dwellers that form small groups or colonies (up to several hundreds) (IUCN 2021). As previously reviewed in Hassanin et al. (2021a,b), there is strong evidence that the genus *Rhinolophus* consitutes the reservoir host in which sarbecoviruses have evolved for centuries. The sarbecoviruses are thought to circulate among bat populations of the main reservoir host species, but other bat species may be occasionally or regularly infected. Importantly, the six groups of bat *Sarbecovirus* genomes showing different SNCs have distinct geographic distributions: China and several bordering countries for *SCoVrC*; southern Yunnan and mainland Southeast Asia for *SCoV2rC*; Yunnan for *YunSar*; Japan for Rc-o319; Bulgaria for RbBM48-31; and Kenya for RspKY72. This suggests that the six groups of viruses have evolved in different *Rhinolophus* reservoirs, each of them being potentially represented by several ecologically related species. Out of Asia, there are two groups of sarbecoviruses, each of them known from a unique virus (Figure 5): RbBM48-31 isolated from *Rhinolophus blasii* in Bulgaria (southeastern Europe) (Drexler et al. 2010), and RspKY72 from Kenya (East Africa), for which the *Rhinolophus* species was not identified in Tao and Tong (2019). In Asia, there are four groups of sarbecoviruses: Rc-o319, *SCoVrC*, *SCoV2rC*, and *YunSar* (Figure 5). The Rc-o319 virus was recently discovered in Japan using fecal samples from *Rhinolophus cornutus* (Murakami et al. 2020), a species endemic to Japanese islands (Burgin et al. 2020). The high nucleotide divergence between Rc-o319 and other sarbecoviruses (between 20% and 26%) supports its evolution in allopatry due to the insular isolation of its bat reservoir. It must be however noted that the species *Rhinolophus nippon* (which is still treated as a subspecies of *Rhinolophus ferrumequinum* in the classification of IUCN, but not in that of Burgin et al. 2020) is distributed on both sides of the Sea of Japan, suggesting that some sarbecoviruses may have occasionally dispersed through long-distance flights between bat populations from the Korean Peninsula and Japan. For *SCoVrC*, many genomic sequences were published during the two last decades because sarbecoviruses have been actively sought in all Chinese provinces after the 2002-2004 SARS epidemic. Although a few viruses were detected in bat genera other than *Rhinolophus*, such as *Aselliscus* (Hu et al. 2017) or *Chaerephon* (Yang et al. 2013), the great majority of *SCoVrC* were isolated from *Rhinolophus* species, and most of them were found in *Rhinolophus sinicus*. The available data suggest therefore that *Rhinolophus sinicus* could be the main reservoir species for *SCoVrC*. In support of this hypothesis, the distribution of *Rhinolophus sinicus* (Burgin et al. 2020; IUCN 2021) fits well with the ecological niche recently inferred for *SCoVrC* (Hassanin et al. 2021b). It appears much more difficult to determine the main reservoir host species (if any) for *SCoV2rC*. Indeed, the five currently known *SCoV2rCs* were identified in five distinct species of Chiroptera: *Rhinolophus affinis*, *Rhinolophus malayanus*, and *Rhinolophus pusillus* in Yunnan (Zhou H. et al. 2020, 2021; Zhou P. et al. 2020), *Rhinolophus acuminatus* in eastern Thailand (Wacharapluesadee et al. 2021) and *Rhinolophus shameli* in northern Cambodia (Hul et al. 2021). Two of these species, *Rhinolophus affinis* and *Rhinolophus pusillus*, are assumed to be largely distributed in China and Southeast Asia (IUCN 2021), but they belong to two different species complexes in which the taxonomy is confused and needs to be clarified (Wu et al. 2012; Soisook et al. 2016; Srinivasulu et al. 2019; Mao and Rossiter 2020). The three other species, *Rhinolophus acuminatus*, *Rhinolophus malayanus*, and *Rhinolophus shameli* are endemic to Southeast Asia (Burgin et al. 2020; IUCN 2021), although the distribution of *Rhinolophus malayanus* has been recently extended to the Yunnan province (Liang et al. 2020). As a consequence, the ecological niche predicted for *SCoV2rC* was found to be different from that of *SCoVrC* (Hassanin et al. 2021b): it includes southern Yunnan and several regions of Laos, Vietnam, Cambodia, Myanmar and Thailand. The two *YunSar* viruses here analysed (RmYN05 and RstYN04) were recently described from two different bat species, *Rhinolophus malayanus* and *Rhinolophus stheno*, collected between May 2019 and November 2020 in Mengla county, Yunnan province (Zhou et al. 2021). The *YunSar* genomes are divergent from other *Sarbecovirus* genomes (between 23% and 27%) and their SNC revealed extremes values for many variables (2/12 in Table 2; 12/32 in Table 3; 19/59 in Table 4). More recently, eight *YunSar* viruses showing 98% of genomic identity with RmYN05 and RstYN04 have been described from *Rhinolophus affinis* and *Rhinolophus stheno* collected in May 2015 in Mojiang County, Yunnan province (Guo et al. 2021). Although current data indicate that the *YunSar* group is endemic to Yunnan, many regions should be explored to better characterise its geographic distribution, including North East India, nothern Myanmar, nothern Laos and nothern Vietnam.

Biogeographically, the most striking result of this study is that the first dimension of all PCAs of Figures 2-4 allowed to separate temperate *versus* tropical groups of *Sarbecovirus*. Indeed, two latitudinal groups can be separated in Asia (Figure 5): the tropical group contains *SCoV2rC*, for which the ecological niche was inferred to include southern Yunnan and several regions of minaland Southeast Asia (Hassanin et al. 2021b), and *PangSar*, for which the most likely origin is Southeast Asia (Hassanin et al. 2021a); and the temperate group is composed of *SCoVrC*, for which the ecological niche was inferred to contain most southern and eastern provinces of China, as well as the Korean Peninsula, Japan, Taiwan, northeastern India, and northern regions of Myanmar and Vietnam (Hassanin et al. 2021b), Rc-o319, which is virus from Japan, and *YunSar*, which is currently endemic to Yunnan. Similarily, two latitudinal groups can be separated in the western Old World (Figure 5): RspKY72 from Kenya in East Africa *versus* RbBM48-31 from Bulgaria in Europe. I suggest that hibernation of bat reservoirs could explain the SNC differences here observed between *Sarbecovirus* genomes from temperate *versus* tropical latitudes. Indeed, most temperate species of *Rhinolophus* found in China, Europe and Japan have to hibernate in winter (Burgin et al. 2020; IUCN 2021) when insect populations become significantly less abundant. By contrast, *Rhinolophus* species found at intertropical latitudes, i.e., between the Tropics of Capricorn (23°S) and Cancer (23°N), do not hibernate because insects are available all year round. In temperate climates, bat hibernation can impact the SNC of *Sarbecovirus* genomes *via* two possible mechanisms: (i) viral replication can be significantly reduced in hibernating bats, and this may explain the lower winter prevalence of coronavirus in hibernating bats (e.g., Lo et al. 2020 for Korean bats); and (ii) the concentrations of free nucleotides available in the cells of hibernating bats can be strongly modified due to the reduction and remodelling of many metabolic pathways (Andrews 2007).

### Why *SCoV2rC* and *PangSar* show similar but different SNCs?

All PCAs of this study showed that the SNCs of *SCoV2rC* genomes are similar to those found for *PangSar* genomes. Indeed, the two groups share extreme values (highest or lowest percentages) for many variables (highlighted in green or red in Tables 2-4), including the highest percentages of A nucleotide and lowest percentage of G at third codon-positions, the highest percentages of U nucleotide and lowest percentages of C and G nucleotides at four-fold degenerate third codon-positions, as well as the highest percentages of A at two-fold degenerate third codon-positions. As previously discussed, these results suggest higher levels of C=>U and G=>A transitions in the genomes of *SCoV2rC* and *PangSar* than in those of other viral lineages (i.e., RbBM48-31, Rc-o319, RspKY72, *SCoVrC*, and *YunSar*). All these elements suggest that *SCoV2rC* and *PangSar* have orginally evolved in the same bat reservoir, which may have included several ecologically related species of *Rhinolophus* distributed in the ecological niche of *SCoV2rC*, i.e., in southern Yunnan and several regions of mainland Southeast Asia (Hassanin et al. 2021b). This hypothesis implies that pangolins are secondary hosts for sarbecoviruses, which is corroborated by the diversity of *Sarbecovirus* lineages found in *Rhinolophus* species (*SCoVrC*, *SCoV2rC*, RbBM48-31, Rc-o319, RspKY72, and *YunSar*) and by the internal placement of the two divergent pangolin viruses in the phylogenetic trees (Figure 1; Zhou H. et al. 2020, 2021; Zhou P. et al. 2020; Hul et al. 2021; Wacharapluesadee et al. 2021).

Despite their high SNC similarities, *SCoV2rC* and *PangSar* genomes appeared in two different clusters in all PCAs (Figures 2-4). In addition, the two groups exhibit special features. On the one hand, the six *SCoV2rC* genomes show the highest percentages of U nucleotide and lowest percentages of C nucleotide at third codon-positions (Table 2). On the other hand, the two *PangSar* genomes (MjGuangdong and MjGuangxi) exhibit the highest percentage of A nucleotide at third codon-positions (Table 2). Importantly, none of the phylogenetic analyses of Figure 1 supported the monophyly of *PangSar*. Based on the whole genomic alignment (Figure 1A), MjGuangdong appeared as more closely related to *SCoV2rC* (BP = 83) than MjGuangxi. The MjGuangdong genome shares between 88% and 90% of identity with *SCoV2rC* genomes, but only 85% with the MjGuangxi genome. By contrast, MjGuangdong was related to *YunSar* in the RdRp gene tree (BP = 99; Figure 1C), and appeared as the sister-group of RshSTT200, MjGuangxi, and SARS-CoV-2 + RaTG13 in the Spike gene tree (BP = 89; Figure 1D). Taken together, all these results suggest that the similar SNCs observed in the two *PangSar* genomes (MjGuangdong and MjGuangxi) were not inherited from a common ancestor, but have been rather acquired by convergence, most probably as a consequence of the shift from their original bat reservoir(s) to Sunda pangolins. As discussed previously, the replication process is dependent of the pool of free nucleotides available in the host cell. As a consequence, variations in the concentrations of ATP, CTP, GTP and UTP among mammalian species (Traut 1994) may have imposed different mutational pressures in case of viral host-shift to a new mammalian species, i.e., from bats to pangolins. In this regard, it is important to note that the human SARS-CoV-2 genome (NC_045512, patient admitted to the Central Hospital of Wuhan on 26 December 2019; Wu et al. 2021) was never grouped to *PangSar* genomes in the PCAs based on SNC (Figures 2-4), whereas it was always closely associated with the five bat *SCoV2rC* genomes. The results support therefore that SARS-CoV-2 emerged directly from a bat virus, without pangolin intermediate host.

## Acknowledgments

This research was funded by the AAP RA-COVID-19, grant number ANR-21-CO12-0002.

## Notes

### Competing Interest Statement

The authors have declared no competing interest.

